# Refractive index modulation by ultraviolet absorption of canonical amino acids for *in vivo* optical transparency

**DOI:** 10.64898/2026.08.22.746458

**Authors:** Su Zhao, Zhongyu Liu, Ling-Yi Zhang, Xuandi Hou, Ani Baghdasaryan, Han Cui, Victoria Crunkleton, Carl H. C. Keck, Dominic Myung, Tianming M. Yang, Kerriann M. Casey, Patrice M. Witschen, Guosong Hong

## Abstract

The inherent opacity of most mammalian tissues limits deep-tissue optical imaging and light delivery. In contrast, the natural transparency of certain species and ocular tissues has been hypothesized to involve proteins with unusually high refractive indices. Here, we systematically analyze the ultraviolet absorption and visible-range refractive index modulation of canonical amino acids to identify key contributors to high-refractive index proteins. We identify arginine hydrochloride as a leading candidate, combining strong ultraviolet absorption, efficient refractive index modulation, physiological pH, and biocompatibility. These properties are validated through successful achievement of optical transparency in both *ex vivo* and *in vivo* tissues. Our findings establish a foundation for using abundant endogenous biomolecules to achieve in vivo tissue transparency and suggest a strategy for engineering proteins enriched in high-performing amino acids to enable efficient, biocompatible tissue clearing.

## Introduction

Optical methods are widely used in biology and medicine, ranging from fluorescence microscopy to optical neuromodulation, fluorescence-guided surgery, and photodynamic therapy.^1–8^ However, the inherent scattering of photons in tissues severely restricts the penetration depth of visible light, often to less than 1 mm, thus limiting noninvasive deep-tissue imaging of biological structures and activity, as well as light delivery for modulating biological function.^9^ Light scattering arises from differences in refractive indices (RIs) between low-RI aqueous components (e.g., the intracellular and extracellular fluid) and high-RI lipid/protein components (e.g., plasma/organelle membranes and collagen fibers in the extracellular matrix).^10^ Tissue clearing technologies can minimize scattering by reducing this RI mismatch. However, conventional tissue clearing methods are incompatible with live tissues as they are often toxic and involve removal of water or lipids, both of which are essential for sustaining life.^11^ Therefore, deep-tissue light delivery and imaging in live mammals still rely on invasive approaches, such as surgically removing the overlying tissue, installation of intravital windows, and invasive insertion of optical fibers and microendoscopes.^12,13^

In stark contrast to the intrinsic opacity of mammalian tissues, some animals, such as zebrafish larvae^14^ and glass frogs^15^, exhibit natural transparency, making them valuable research organisms. It has been hypothesized that reduced RI heterogeneity in these organisms contributes to suppressed optical scattering, potentially through the endogenous production of water-soluble high-RI molecules that act as intrinsic optical clearing agents. Examples of endogenous proteins with high RI and excellent water solubility include antifreeze proteins and crystallins.^16^ Crystallins, in particular, are responsible for the high RI and optical transparency of the lens, one of the few inherently transparent tissues in mammals.^17^ Together, these observations suggest that nature exploits the enrichment of high-RI molecular components as a strategy to minimize optical scattering in specialized transparent tissues.

The ability of biological systems to produce some proteins with higher RIs than others prompts the question of why this occurs. Although there is a broad consensus that the RI increment of most proteins lies within a narrow range of ∼0.185 mL/g (±2%), some proteins represent high-end outliers.^18^ For example, crystallins have been reported to deviate substantially from this range, with RI increments of 0.1930 ± 0.0055 mL/g and the highest-RI crystallins approaching 0.199 mL/g. At a concentration of 1 g/mL, such crystallins would increase the RI of an aqueous solution by ∼0.199, corresponding to a solution RI of ∼1.53. Similarly, fatty acid hydroxylases constitute another class of high-RI proteins (∼0.197 mL/g), likely due to their elevated aromatic amino acid content. However, unlike crystallins, these proteins are predominantly membrane-associated and therefore lack high water solubility. These observations suggest that a protein’s RI depends on its specific amino-acid composition, which determines its contribution to the visible RI through ultraviolet absorption, in accordance with the Kramers-Kronig relations (**Fig. 1**).

**Fig. 1.**
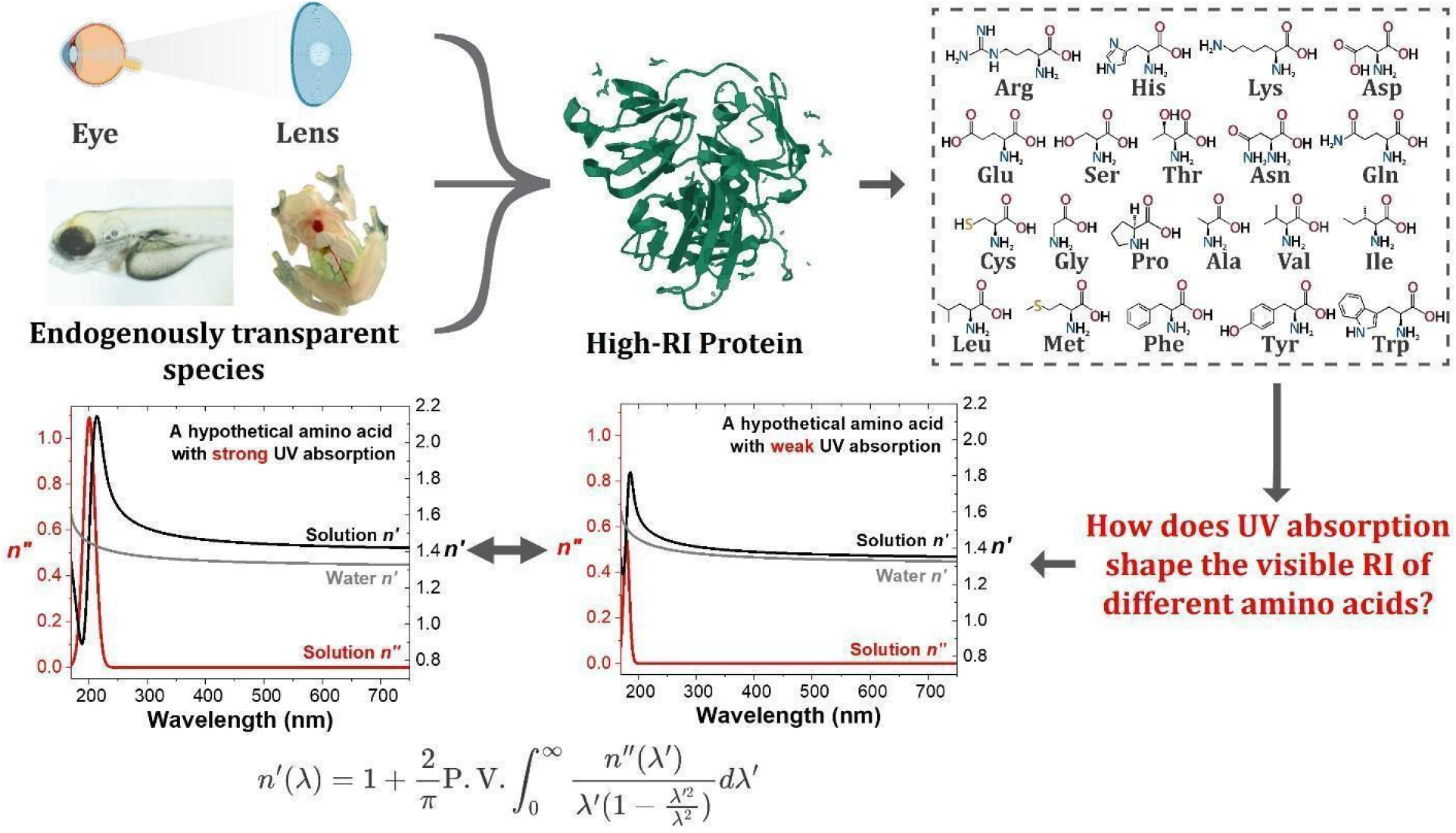
The high refractive indices (RIs) of endogenous proteins in biological tissues motivate this study to identify amino acids with high RI based on their strong ultraviolet absorption, in accordance with the Kramers-Kronig relations (equation shown near the end of the figure).

This work is inspired by inherently high-RI proteins, whose elevated indices are linked to strong ultraviolet absorption. Motivated by this insight, we systematically screened all twenty canonical amino acids to evaluate their ultraviolet absorption characteristics and their ability to modulate the RI of water upon dissolution. We found that the three aromatic amino acids – tryptophan (Trp), tyrosine (Tyr), and phenylalanine (Phe) – exhibit the strongest ultraviolet absorption, consistent with their reported high RI increments as predicted by the Kramers-Kronig relations.^19,20^ Beyond aromatic residues, several amino acids with delocalized electrons arising from conjugation or heteroatoms, such as arginine (Arg) and cysteine (Cys), also display substantial ultraviolet absorption and high RI increments. Notably, these amino acids are enriched in canonical high-RI proteins, including crystallins.^18,21^ Importantly, Arg offers practical advantages due to its high aqueous solubility in the salt form (i.e., Arg·HCl), enabling significant RI modulation at physiologically neutral pH – an advantage over its aromatic counterparts. Importantly, we demonstrate that Arg·HCl can induce optical transparency in excised *ex vivo* mouse skin as well as in the *in vivo* mouse abdomen with favorable local and systemic biocompatibility. Collectively, this work elucidates fundamental biophysical principles underlying intrinsically high-RI proteins and introduces a new class of biologically derived, biocompatible clearing agents for achieving *in vivo* tissue transparency.

## Results and Discussion

### Ultraviolet absorption of amino acid solutions

Because the high RIs of proteins – such as crystallins in the ocular lens and proteins found in inherently transparent animals (**Fig. 1**) – originate from their constituent amino acids and the intermolecular interactions, we first sought to address a fundamental question: which classes of amino acids contribute most significantly to the elevated RI of these proteins? The real part of the RI, *n′*, is causally linked to its imaginary component, *n“*, through the Kramers-Kronig relations, as described below:

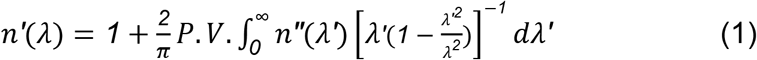

where P.V. is the Cauchy principal value of the integral, and both *n′* and *n“* are functions of wavelength *λ*. Here, the imaginary component *n“*(*λ*) is directly related to optical absorption according to the following equation:

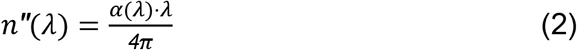

where *α*(*λ*) is the wavelength-dependent absorption coefficient of the material. Therefore, by measuring the ultraviolet absorption spectra of all twenty canonical amino acids, we can infer the origins of their differing real RIs in the visible spectrum through Eq. (1).

We measured the ultraviolet absorption spectra of aqueous solutions of all twenty canonical amino acids and plotted their molar absorption coefficients, *ε*, in **Fig. 2**. The molar absorption coefficients, *ε*, is directly proportional to the absorption coefficient, *α*, as follows:

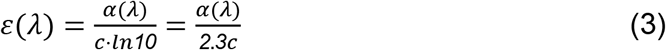

**Fig. 2.**
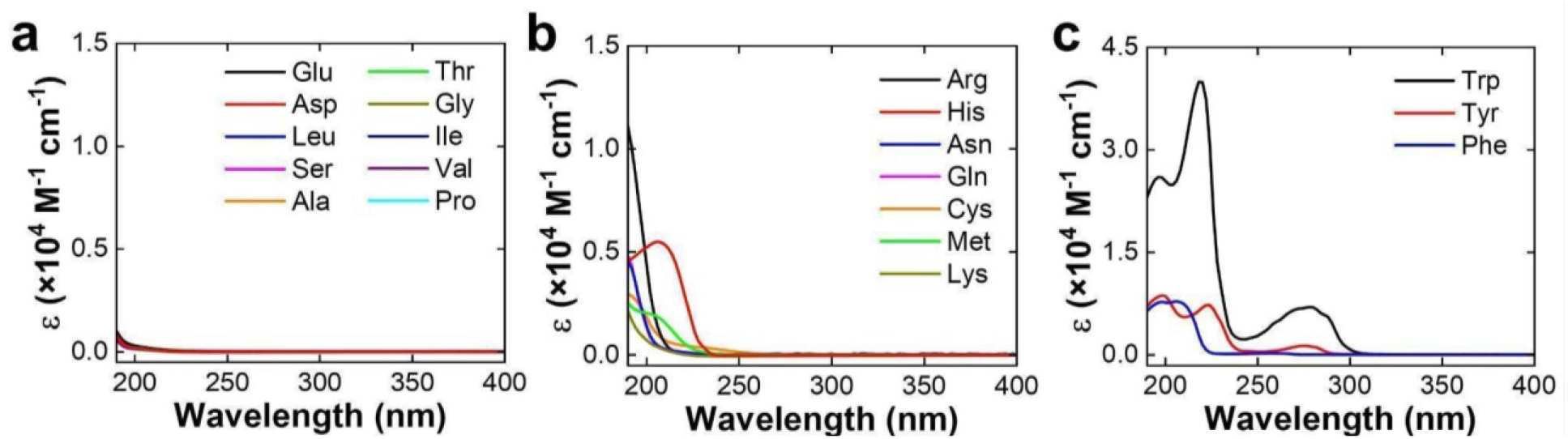
Ultraviolet absorption spectra of (**a**) weakly absorbing amino acids due to their low absorbance in the plotted spectral region, (**b**) moderately absorbing amino acids with overlapping spectra for Gln and Asn, and (**c**) strongly absorbing amino acids. Spectra were measured at 0.5 mg/mL for all solutions.

where *c* represents the molar concentration, in mol/L or M, of the amino acid in the solution. To ensure that all amino acids were fully dissolved and measured under identical conditions, each solution was prepared at a concentration of 0.5 mg/mL. The measured absorbance was then normalized using the molecular weight of each amino acid to obtain *ε*. We note that at higher concentrations, the molar absorption coefficient for the amino acids may deviate substantially from Beer-Lambert behavior.^22,23^ As a result, we chose a dilute concentration of 0.5 mg/mL to perform these absorption measurements in **Fig. 2**.

The ultraviolet molar absorption spectra, *ε*(*λ*), of the twenty canonical amino acids reveal several notable trends. Specifically, ten of the twenty amino acids exhibit extremely weak absorption across the measured spectral range from 190 to 400 nm (**Fig. 2a**). A common feature of these amino acids is the absence of delocalized π systems or highly polarizable heteroatoms (e.g., sulfur) in their side chains. Members of this group predominantly possess simple aliphatic side chains (Leu, Ala, Gly, Ile, Val, and Pro) or side chains containing oxygen-based functional groups only, such as carboxyl or hydroxyl moieties (Glu, Asp, Ser, and Thr). These aliphatic, carboxyl, and hydroxyl groups are known to exhibit low electronic polarizability due to the small atomic radii of their constituent atoms (C, H, and O) and the limited extent of electron delocalization.^24^ As a result, the resonant absorption wavelengths of these functional groups typically lie below 190 nm, outside our measurement window and far from the visible spectrum. Consequently, their contribution to RI modulation in the visible range is minimal, as predicted by Eq. (1). This follows from the 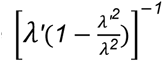 weighting in the Kramers-Kronig relation, which indicates that absorption features increasingly distant from the wavelength of interest exert a diminishing influence on the real RI.

In contrast to the first group of ten amino acids, which exhibit negligible absorption features in the 190-400 nm range, a second group comprising seven amino acids displays stronger and red-shifted absorption profiles across this spectral window (**Fig. 2b**). A defining characteristic of this group is the presence of delocalized π systems or highly polarizable heteroatoms (e.g., sulfur) in their side chains, which act as ultraviolet chromophores. For example, histidine (His) contains an imidazole ring – an aromatic heterocycle with delocalized π electrons – in its side chain. Arginine (Arg) contains a guanidinium group, which is known to exhibit “Y-aromaticity” with stably delocalized π electrons despite its acyclic structure.^25^ The remaining amino acids in this group incorporate polarizable heteroatoms, such as sulfur (Cys and Met) or nitrogen (Asn, Gln, and Lys), within their side chains.

Both delocalized π systems and highly polarizable heteroatoms reduce the effective electronic resonance frequency, *ω_0_*, leading to red-shifted absorption peaks and enhanced absorption strength, as captured by the Lorentz oscillator model:

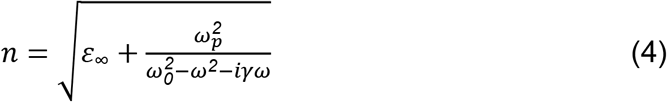

where a clear inverse dependence of the complex refractive index, *n*, on *ω_0_*can be observed. As a result, these amino acids are expected to exhibit higher RI increments in the visible spectrum. This expectation is supported by the elevated RIs of arginine-rich and methionine-rich proteins such as crystallins. Notably, human γ-crystallins are enriched in arginine, whereas fish γ-crystallins exhibit a higher methionine content relative to the proteome average.^26^ These findings reflect convergent evolutionary strategies for achieving the high RI required for lens function (e.g., refractive focusing) through enrichment of ultraviolet-absorbing amino acid residues.

Finally, a third group comprising three amino acids – tryptophan (Trp), tyrosine (Tyr), and phenylalanine (Phe) – exhibits exceptionally strong absorption across the 190-400 nm range (**Fig. 2c**). A defining feature of these amino acids is the presence of highly delocalized π systems, such as benzene and indole rings, in their side chains. Notably, tryptophan displays a peak molar absorption coefficient of ∼4×10^4^ M^−1^cm^−1^, making it the most strongly absorbing canonical amino acid due to the strong π-π* transitions of its indolyl group. Consistent with their strong ultraviolet absorption, these amino acids also exhibit exceptionally high RI increments, as reported previously.^18^ For example, tryptophan reaches an RI increment of 0.277 mL/g, substantially higher than that of minimally absorbing amino acids in the first group (e.g., 0.170 mL/g for serine). These observations help explain why proteins enriched in aromatic amino acids – such as fatty acid hydroxylases, where aromatic residues are thought to facilitate substrate binding – often exhibit unusually high RI increments.^18,27^

### Refractometry and ellipsometry of amino acid solutions

We next sought to determine whether the ultraviolet absorption of these twenty amino acids correlates with RI increments in the visible spectrum. To this end, we measured the RI of aqueous solutions of the same twenty amino acids at a high concentration of 3 M. Both a digital refractometer and a spectroscopic ellipsometer were employed to measure the real refractive index at 589 nm and the complex refractive-index spectrum over the 250-750 nm range, respectively.

This high concentration of 3 M amino acids was chosen for two reasons. First, unlike the ultraviolet absorption measurements in **Fig. 2**, which were performed at low concentrations to minimize errors associated with transmission-based spectroscopy, ellipsometry requires a sufficiently large RI contrast relative to pure water to yield reliable measurements. Second, the concentration of high-RI proteins in certain inherently transparent tissues – such as crystallins in the ocular lens – can exceed 500 mg/mL, corresponding to an effective amino-acid residue concentration of approximately 3 M.^26,28^ Thus, using a 3 M solution allows us to accurately quantify RI increments while also approximating the physiologically relevant concentration regime of high-RI proteins *in vivo*.

We prepared 3 M amino acid solutions in their original form, as well as in protonated (with equimolar HCl) and deprotonated (with equimolar NaOH) forms. Measurements of their solubility, refractive indices, and pH revealed several key findings (**Table 1**). First, among the twenty canonical amino acids tested, only lysine, serine, glycine, and proline were soluble at 3 M in their original forms. Second, many amino acids possess isoelectric points (pI) far from neutral pH (e.g., arginine has a pI of 10.76), making them difficult to dissolve at 3 M in their zwitterionic forms. Indeed, in proteins under physiological conditions, the ionizable side chains of these amino acids are predominantly protonated (for basic residues) or deprotonated (for acidic residues). Accordingly, we prepared 3 M solutions of these amino acids in their protonated and/or deprotonated forms as appropriate (see **Materials and Methods**). Notably, protonated arginine (Arg·HCl) and protonated lysine (Lys·HCl) both achieve relatively high RI values (1.450 and 1.433, respectively) at near-physiological pH (7.73 and 7.67). These elevated RI values are consistent with the moderately strong UV absorption of Arg and Lys (**Fig. 2b**).

**Table 1.** Refractive index and pH of the 20 amino acids in their original, protonated, and deprotonated forms. The concentration is 3 M for all soluble forms reported in the table.

| Amino acid | RI, original | pH, original | RI, protonated | pH, protonated | RI, deprotonated | pH, deprotonated |
| --- | --- | --- | --- | --- | --- | --- |
| Arginine (Arg) | Insoluble | Insoluble | 1.450 | 7.73 | Insoluble | Insoluble |
| Histidine (His) | Insoluble | Insoluble | Insoluble | Insoluble | 1.430 | 12.14 |
| Lysine (Lys) | 1.412 | 10.46 | 1.433 | 7.67 | 1.417 | 13.56 |
| Aspartic acid (Asp) | Insoluble | Insoluble | 1.400 | 0.24 | 1.398 | 9.52 |
| Glutamic acid (Glu) | Insoluble | Insoluble | Insoluble | Insoluble | 1.415 | 7.07 |
| Serine (Ser) | 1.384 | 6.05 | 1.398 | 0.34 | 1.390 | 11.90 |
| Threonine (Thr) | Insoluble | Insoluble | 1.404 | 0.49 | 1.397 | 12.23 |
| Asparagine (Asn) | Insoluble | Insoluble | 1.414 | 0.61 | 1.407 | 11.62 |
| Glutamine (Gln) | Insoluble | Insoluble | Insoluble | Insoluble | 1.414 | 11.89 |
| Cysteine (Cys) | Insoluble | Insoluble | 1.416 | 0.61 | 1.430 | 9.72 |
| Glycine (Gly) | 1.371 | 6.54 | 1.385 | 0.71 | 1.376 | 12.36 |
| Proline (Pro) | 1.392 | 7.06 | 1.405 | 0.56 | 1.400 | 13.35 |
| Alanine (Ala) | Insoluble | Insoluble | 1.390 | 0.70 | 1.383 | 12.47 |
| Valine (Val) | Insoluble | Insoluble | Insoluble | Insoluble | 1.402 | 12.37 |
| Isoleucine (Ile) | Insoluble | Insoluble | Insoluble | Insoluble | 1.406 | 12.44 |
| Leucine (Leu) | Insoluble | Insoluble | Insoluble | Insoluble | 1.402 | 12.20 |
| Methionine (Met) | Insoluble | Insoluble | 1.431 | 0.71 | 1.425 | 12.19 |
| Phenylalanine (Phe) | Insoluble | Insoluble | Insoluble | Insoluble | 1.443 | 11.68 |
| Tryptophan (Trp) | Insoluble | Insoluble | Insoluble | Insoluble | 1.488 | 11.80 |
| Tyrosine (Tyr) | Insoluble | Insoluble | Insoluble | Insoluble | Insoluble | Insoluble |

Third, deprotonated phenylalanine (Phe·NaOH) and deprotonated tryptophan (Trp·NaOH) achieve some of the highest RI values observed in this study (1.443 and 1.488, respectively), consistent with their strong UV absorption (**Fig. 2c**). However, their extremely basic pH values (approaching 12) in these deprotonated forms make them unsuitable in biologically relevant environments. We hypothesize that their solubility in NaOH arises not from a low pI, but from the net negative charge of the deprotonated species, which electrostatically stabilizes the anions in water. Finally, it is worth noting that tyrosine was the only amino acid in our study that did not fully dissolve at 3 M in any form tested. As a result, its RI increment at the benchmark concentration of 3 M could not be reliably evaluated despite its strong UV absorption (**Fig. 2c**).

In addition to refractometry, the complex RI spectra of the amino acid solutions over the 250-750 nm range were measured using spectroscopic ellipsometry (**Figs. S1-S3**). The ellipsometry results are in good agreement with those obtained from the refractometer. Importantly, all amino acids exhibit a decrease in the real part of the RI, *n′*, with increasing wavelength beyond 300 nm. This trend represents a characteristic dispersion behavior arising from resonant absorption below 300 nm. Furthermore, this dispersion eventually reaches a plateau in the visible spectrum, although the plateau value of *n′* varies substantially among different amino acids. For example, tryptophan in its deprotonated form exhibits the highest visible RI (*n′* = 1.48) even after reaching the plateau. In contrast, glycine in its original form shows the lowest visible RI (*n′* = 1.37). These results reveal that highly polarizable amino acids, such as tryptophan, can elevate the baseline RI of the surrounding medium (e.g., water) by nearly fourfold compared with those composed of amino acids lacking polarizable side chains, such as glycine. This variation closely tracks the strength of ultraviolet absorption and can be attributed to differences in amino acid side-chain chemistry, as shown in **Fig. 2**.

The varying abilities of different amino acids to increase the baseline RI of water, as revealed in **Table 1** and **Figs. S1-S3**, represent not only a scientific discovery but also an engineering principle for identifying potential new clearing agents, either as individual amino acids in their molecular form or as derived peptides. For practical tissue-clearing applications, two key considerations must be taken into account: the RI increment in the visible spectrum and the pH at which specific amino acids achieve such RI increments. To facilitate visualization of these two factors, we plot all amino acids that are soluble at 3 M, in their original, protonated, and deprotonated forms, according to their RI and solution pH (**Fig. 3**). This chart clearly highlights amino acids that satisfy both RI and pH requirements, located in the bottom-right region of the graph, such as Arg·HCl. Specifically, Arg·HCl achieves a high RI of 1.45 while maintaining a near-physiological pH of 7.7 in a 3 M aqueous solution (**Fig. 3** and **Table 1**). Interestingly, this observation is also consistent with the enrichment of protonated arginine residues in endogenously high-RI proteins, such as crystallins, across different species.^26^

**Fig. 3.**
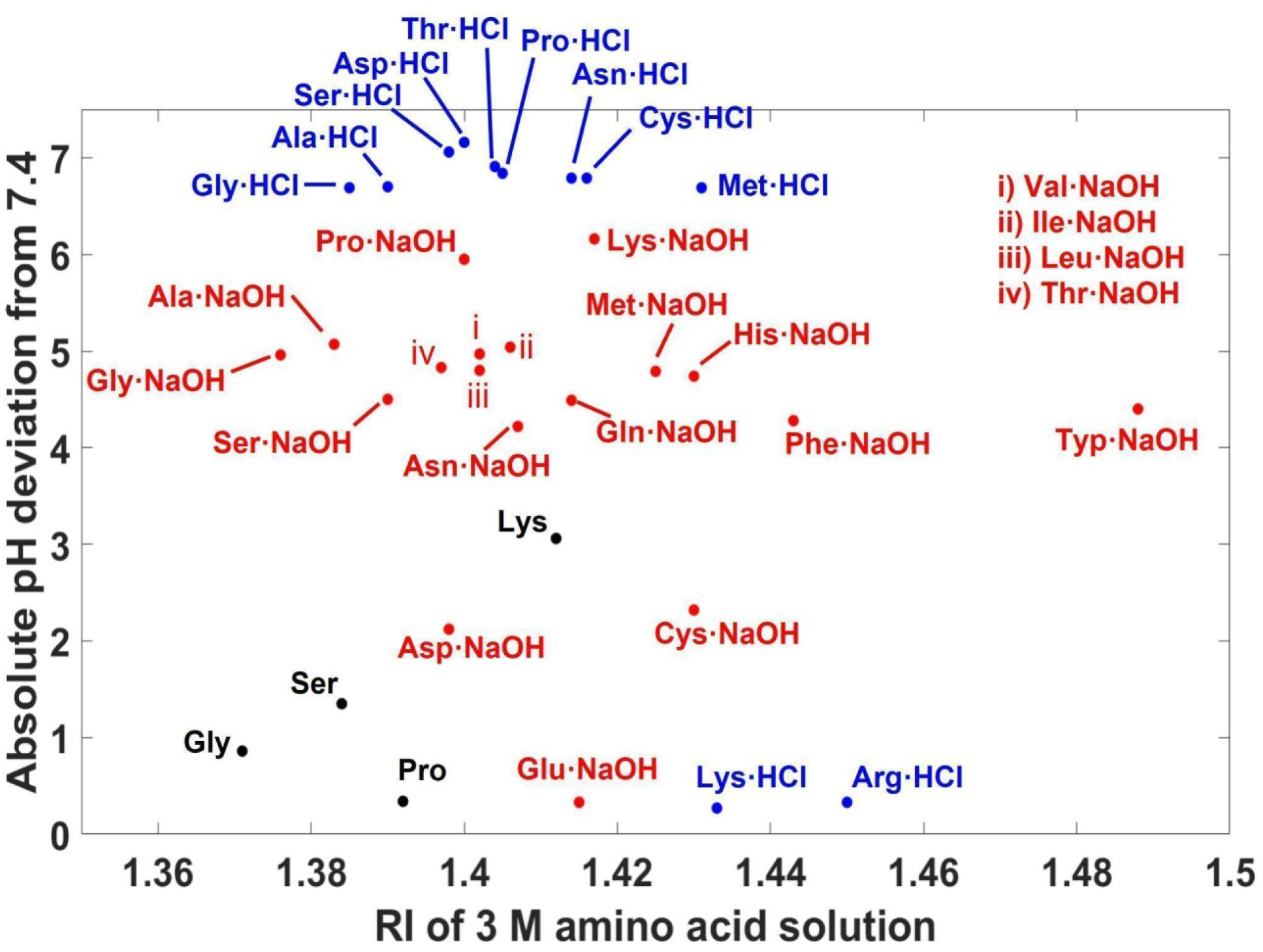
Amino acids, in their original (black), protonated (blue), and deprotonated (red) forms (where applicable), plotted according to their RI and deviation from the physiological pH of 7.4. This figure is plotted based on the data in **Table 1**.

### Optical characterizations and tissue-clearing ability of arginine hydrochloride

The optical properties of the top candidate identified in our screening (**Fig. 3**), Arg·HCl, were further examined. As shown in **Fig. 4a**, Arg·HCl solutions exhibit near-100% transmission beyond 300 nm and across the visible spectrum. This lack of absorption in the visible range is attributed to its absorption being confined to wavelengths below 250 nm (**Fig. 4b**), with absorption scaling linearly with concentration in accordance with the Beer-Lambert law (**Fig. 4c**). In addition to absorption, we measured the RI of Arg·HCl solutions at 589 nm as a function of concentration, revealing a similarly linear relationship (**Fig. 4d**). This linearity further suggests the absence of significant intermolecular interactions among arginine molecules in solution that would otherwise lead to nonlinear enhancement or suppression of absorption and RI at higher concentrations.

**Fig. 4.**
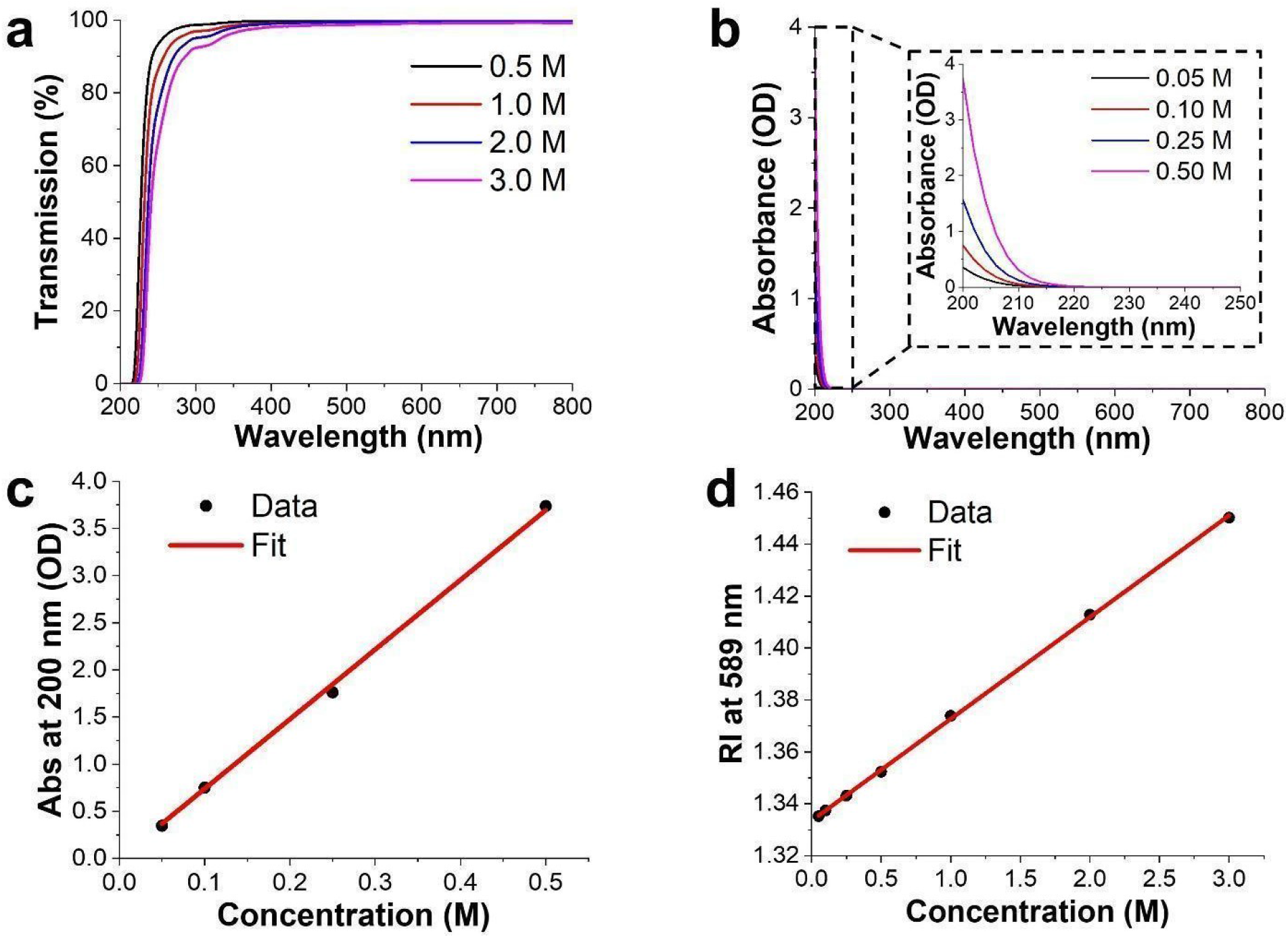
Optical characterizations of Arg·HCl solutions. (**a**) Transmission spectra of Arg·HCl solutions of different concentrations through a 1-mm optical path length in the 200-800 nm range. The small depression at ∼310 nm emerges at high concentrations and is likely attributable to aggregation-induced excited states arising from intermolecular interactions among arginine molecules.^55^ (**b**) Absorbance spectra of Arg·HCl solutions of different concentrations through a 10-μm optical path length in the 200-800 nm range. The inset shows a closeup view of the spectra in the 200-250 nm window that corresponds to the dashed box. (**c**) Absorbance of Arg·HCl solutions at 200 nm plotted against their concentrations, revealing a linear relationship consistent with the Beer-Lambert law. Linear regression yields a slope of 7.39 M^−1^, which corresponds to an extracted molar absorption coefficient of 7.39×10^3^ M^−1^cm^−1^ for Arg·HCl at 200 nm. (**d**) Refractive index of Arg·HCl solutions at 589 nm plotted against their concentrations, revealing a linear relationship with an extracted molar RI increment of 0.039 M^−1^.

The high RI of Arg·HCl solutions, together with the implication of protonated arginine in endogenous high-RI proteins and inherent tissue transparency, prompted us to evaluate its ability to induce optical transparency in otherwise opaque tissues. We first validated the feasibility of using Arg·HCl solutions to induce optical transparency in excised mouse abdominal skin. Specifically, a 2 M Arg·HCl solution (RI = 1.41, **Fig. 4d**) was used, as this value closely matches the RIs of lipids and collagen, the primary scattering components in skin.^10,29^ Although a recent study suggested an optimal RI of 1.36∼1.37 for live tissue clearing, this conclusion was based on dissociated cells and did not account for extracellular scatterers present in intact tissues.^30^ In contrast, recent work from our lab demonstrates improved RI matching and reduced scattering at RI values up to 1.41.^31^

A piece of freshly dissected mouse skin (approximately 10 mm × 20 mm) was used as an experimental sample, which was treated either with Arg·HCl solution or an osmolarity-matched NaCl solution as the control.^32^ As shown in **Fig. 5**, Arg·HCl induced substantial optical transparency in mouse skin over the course of a few hours, whereas no noticeable improvement was observed with the osmolarity-matched NaCl solution.

**Fig. 5.**
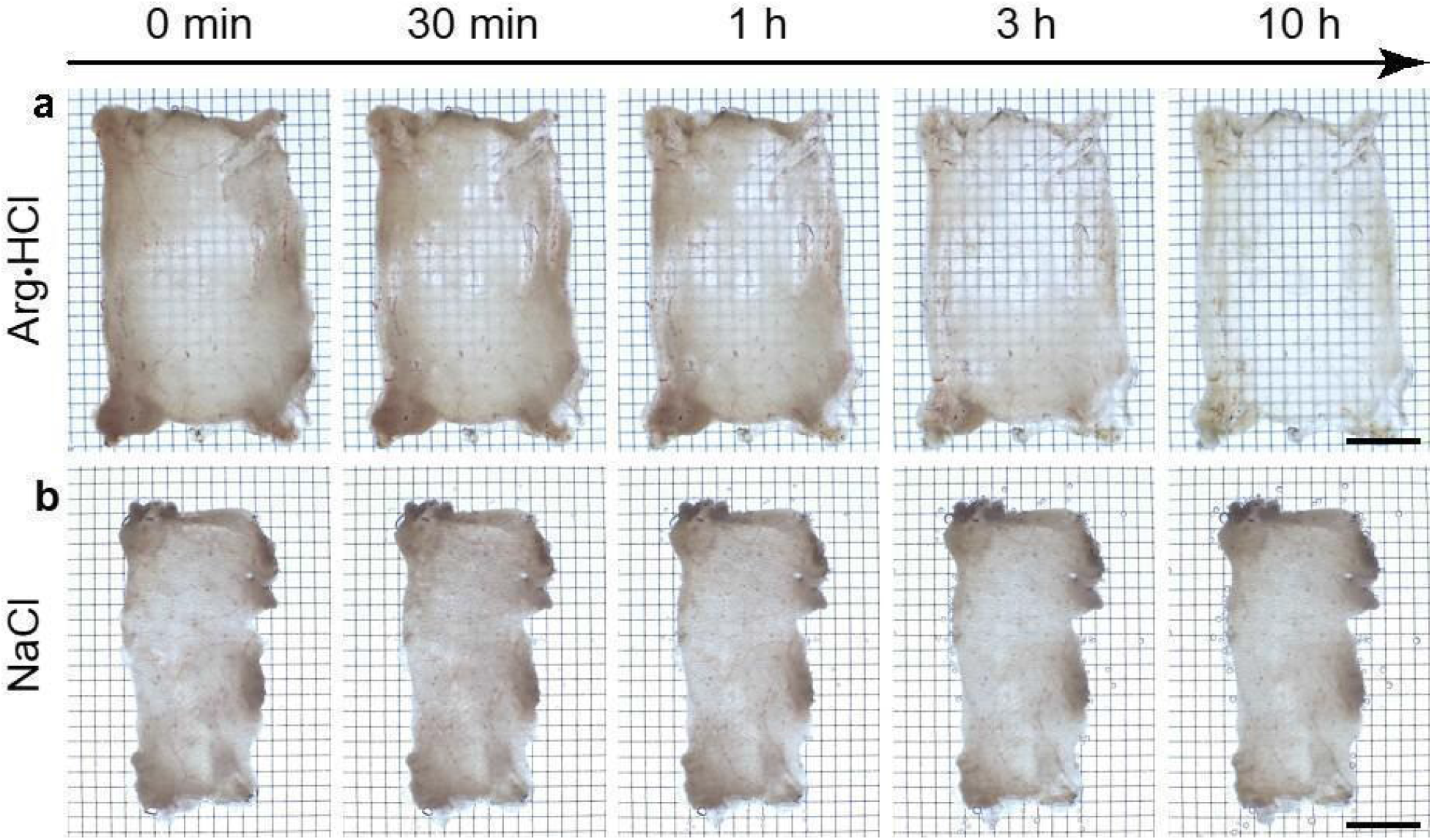
Arginine hydrochloride achieves optical transparency in *ex vivo* mouse skin. Brightfield transmission images of the mouse skin taken at 0 min, 30 min, 1 h, 3 h, and 10 h after soaking in (**a**) 2 M Arg⋅HCl solution and (**b**) osmolarity-matched NaCl solution. Scale bars: 5 mm.

We next sought to demonstrate the ability of amino acid solutions to induce tissue transparency in the abdominal skin of live mice (**Fig. 6a**). Anesthetized C57BL/6 mice aged 4-5 weeks and weighing 11-13 g were used to enable direct visualization of internal organs, as older mice possess greater subcutaneous fat that obscures optical access.^32–34^ Notably, skin from older mice can still be rendered transparent using the *in vivo* clearing approach when combined with imaging modalities such as optical coherence tomography (OCT), which is less susceptible to fat-induced signal attenuation, or when applied to anatomical regions with consistently low subcutaneous fat, such as the scalp, even in mice up to 18 weeks of age.^35,36^ Nair was used to remove abdominal fur and enhance skin permeability by exfoliating the epidermis.

**Fig. 6.**
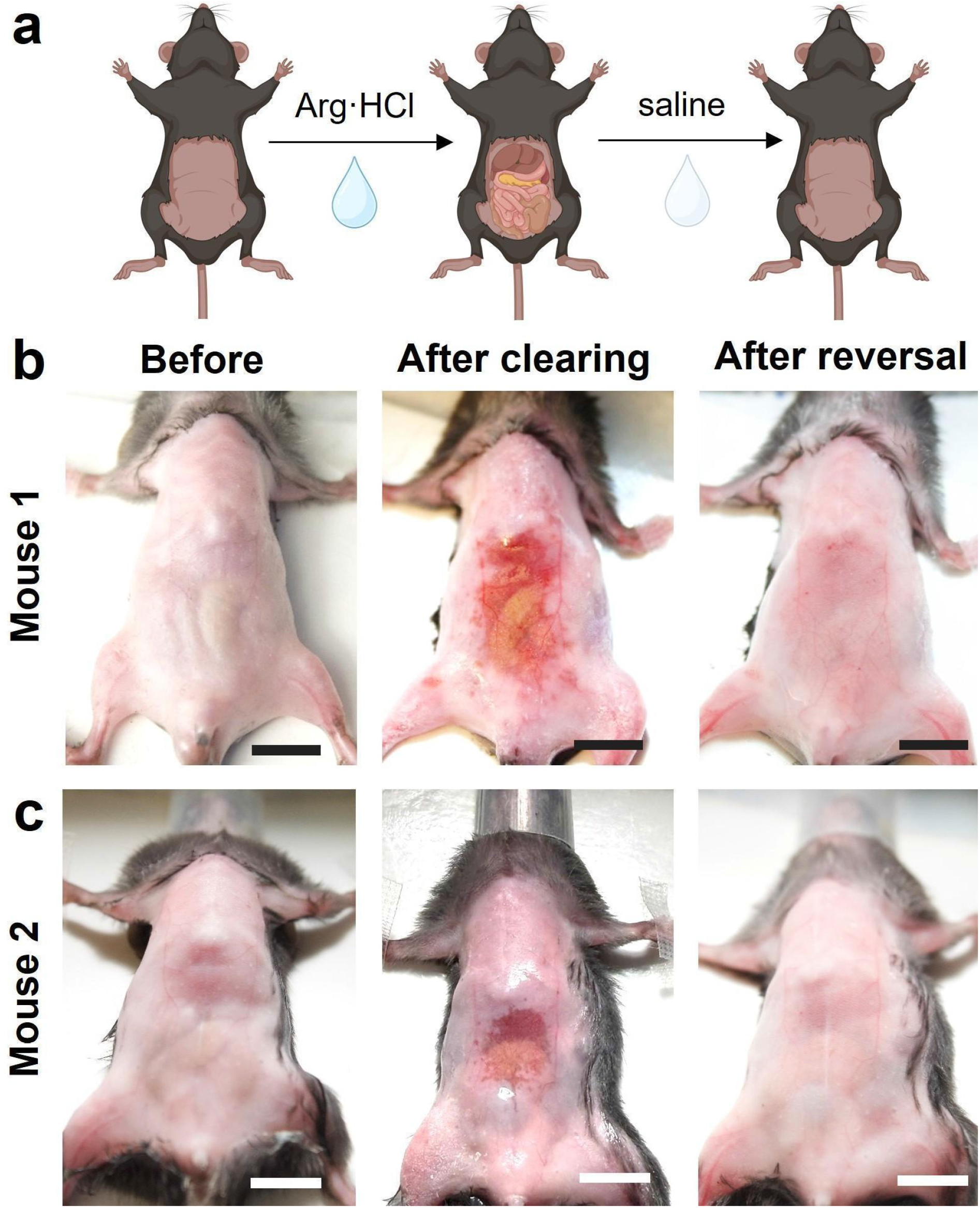
Arginine hydrochloride achieves optical transparency in the abdominal skin of live mice. (**a**) Schematic showing the procedure of achieving and reversing optical transparency in abdominal skin. (**b**,**c**) Representative brightfield images of two mice before treatment (left), after achieving transparency with arginine hydrochloride (middle), and after reversing the transparency effect with saline (right). Scale bars: 1 cm.

The Arg·HCl solution was applied topically using a cotton applicator and massaged into the depilated and exfoliated skin for 10 minutes to facilitate penetration. Compared with pre-treatment images, which exhibit the typical opacity of skin that obscures underlying organs, post-treatment images show that arginine hydrochloride induces substantial transparency in the abdominal wall, enabling direct visualization of internal organs such as the liver and intestines (**Fig. 6b,c**; **Movies S1** and **S2**). The transparency effect is reversible: rinsing with saline restores the skin to its original opacity. Unlike previous reports in which tartrazine-induced transparency resulted in orange-to-red skin coloration,^32,33,35,37–46^ the skin retained its natural appearance, allowing visualization of internal organs, such as the intestines, in their native colors.

### Bioeffects of arginine hydrochloride as an *in vivo* optical clearing agent

We next performed a series of experiments to investigate the chronic effects of arginine-induced tissue transparency. Arginine is a canonical amino acid with a reported median lethal dose (LD50) of 12 g/kg body weight,^47^ making it among the safest compounds for *in vivo* use and a widely used dietary supplement.^48^ However, at the concentrations required to achieve efficient transparency (2 M), the high osmolality may induce transient cellular stress prior to washout. Accordingly, we conducted blood chemistry, complete blood count (CBC), gross skin assessment, and skin and major organ histology analyses to evaluate both the short- and long-term effects of arginine hydrochloride-induced tissue transparency.

Comprehensive blood chemistry and CBC analyses performed at 24 h and 14 days after abdominal optical clearing with Arg·HCl revealed no statistically significant differences in any blood chemistry biomarkers and most CBC parameters between the control (saline) and experimental (Arg·HCl) groups (**Figs. S4 and S5**). A statistically significant increase in total WBC count was observed in control animals compared to experimental animals at day 14 (P = 0.016); however, no significant changes were detected in the major leukocyte subpopulations, suggesting the absence of substantial systemic inflammatory or immune-related toxicity (**Fig. S5**).

In addition, we monitored the gross appearance of the treated abdominal skin for two weeks following optical clearing to assess potential signs of inflammation or necrosis (**Fig. S6**). Serial photographs showed largely normal skin appearance, including smooth texture and healthy pink coloration, during the first two days after Arg·HCl-mediated clearing and reversal (**Fig. S6a-b**). Beginning on Day 3, a small region of apparent ulceration, substantially smaller than the cleared area, became visible and resolved by Day 6 (**Fig. S6c-f**). We hypothesize that the increased susceptibility of the skin to dehydration following removal of the stratum corneum to enhance clearing agent penetration is the primary cause of the minor ulceration observed between Days 3 and 6, as described in our recent report.^32^ Hair regrowth became apparent by day 10 and returned to full abdominal coverage by day 14 (**Fig. S6j-o**). Consistent with these observations, histologic evaluation and scoring revealed that tissues from both saline-treated controls and arginine-treated animals did not differ at both day 1 and day 14 (**Fig. S7&S8**; **Table S1**). Lastly, H&E staining of major organs, including the brain, lungs, heart, liver, spleen, and kidneys, revealed no pathological abnormalities related to Arg·HCl-mediated clearing and reversal (**Fig. S9-S14**). Collectively, these findings demonstrate the favorable local and systemic biocompatibility of the arginine-based clearing formulation following topical administration.

### Conclusion and Outlook

This work represents an effort to learn from endogenous proteins and canonical amino acids with high RIs, while also exploring the potential of the identified RI-contributing amino acids as biocompatible agents for *in vivo* optical clearing. Our findings identify strongly UV-absorbing amino acids as major contributors to RI modulation, consistent with the Kramers-Kronig relations and our previously established framework of dye-induced tissue transparency.^32–34,37,49^ Specifically, we identify arginine hydrochloride as both a major contributor to high-RI proteins and a biocompatible clearing agent, owing to its efficient RI modulation, near-physiological pH, and established biosafety. Using arginine hydrochloride, we achieve optical transparency in excised mouse skin and in the live mouse abdomen across the visible spectrum.

One potential concern with high-concentration amino acid solutions is hyperosmolality, which can induce cellular stress, particularly under prolonged exposure.^30^ Our recent study reveals that iso-osmolality is not a universal requirement for in vivo optical clearing agents. Instead, hyperosmolar clearing solutions can be well tolerated by living cells, depending on the specific cell and tissue types to be rendered transparent, the required duration of transparency, and the cell membrane permeability of the clearing agent.^31,37^ Many amino acids, particularly arginine, have been reported to exhibit membrane permeability.^50–53^ Consequently, even when present in hyperosmolar extracellular solutions, their ability to permeate cell membranes can reduce transmembrane osmotic gradients, alleviating the need for strict physiological osmolality by maintaining conditions closer to isotonic across the cell membrane. Consistent with this finding, our comprehensive chronic biosafety studies indicate minimal local and systemic effects (**Fig. S4-S14** and **Table S1**), despite the effective concentrations substantially exceeding physiological osmolality (**Fig. S15**).

The current need for high nominal concentrations of arginine, and the associated hyperosmolality, arises because arginine exhibits only modest UV absorption on a per-molecule basis. The molar extinction coefficient of arginine at its absorption peak is approximately threefold lower than that of a typical visible dye, such as tartrazine,^33^ as predicted by the Lorentz oscillator model (Eq. 4). This reduction is the necessary trade-off for achieving color-free optical clearing.^37^ Although our analysis identifies several amino acids with stronger UV absorption and greater visible-range RI enhancement, such as Phe, Tyr, and Trp, these amino acids generally exhibit limited aqueous solubility and deviate substantially from physiological pH at the concentrations required for optical clearing (**Figs. 2** and **3**; **Table 1**). A promising strategy for future studies is therefore to design and synthesize oligopeptides, polypeptides, or proteins enriched in arginine together with other high-RI-contributing residues (e.g., Phe, Tyr, and Trp). Such engineered molecules could maximize UV absorption and, consequently, RI enhancement in the visible spectrum, while reducing the required molar concentration and osmolality by incorporating multiple high-RI residues into each macromolecule. At the same time, the strong hydration capacity of arginine may improve the aqueous solubility of otherwise poorly soluble aromatic residues. Notably, this design principle is reflected in the molecular composition of crystallins across diverse species, which combine exceptionally high RIs with excellent water solubility.^26^ More broadly, the insights gained in this study provide a framework for engineering proteins enriched in high-RI-contributing residues, raising the possibility of genetically encoding and overexpressing such proteins to impart intrinsic optical transparency to otherwise opaque tissues.

## Methods

### Chemicals

L-asparagine (≥98%, A0884-25G, batch 0000349152), L-alanine (≥98.5%, A7469-25G, batch 0000356249), L-arginine (≥99.5%, 11009-100G-F, batch BCCM5514), L-cysteine (97%, 168149-25G, batch BCCK4081), L-glutamic acid (≥99%, G1251-100G, batch 0000325266), L-glutamine (≥99%, G8540-25G, batch 0000372175), L-histidine (≥99%, H8000-25G, batch 0000324255), L-isoleucine (99.7%, 8160080025, batch S6015008), L-methionine (≥98%, M9625-25G, batch 0000317863), L-proline (≥99%, P0380-100G, batch SLBW7100), L-phenylalanine (≥98.5%, P5482-25G, batch 0000380797), L-serine (≥98.5%, S4311-25G, batch 0000392173), L-threonine (≥98%, T8625-25G, batch SLCR3495), L-valine (≥98.5%, V0513-25G, batch 0000345519), L-lysine (≥98%, L5501-25G, batch BCCP0561), L-aspartic acid (≥99%, A8949-25G, batch BCCJ9947), L-tryptophan (≥98%, T0254-25G, batch WXBF6780V), L-tyrosine (≥99%, T8566-25G, batch 0000511048), L-leucine (≥98%, L8000-25G, batch BCCN4277), and glycine (≥99%, G7126-100G, batch 0000474751) were purchased from Sigma-Aldrich Inc (Burlington, Massachusetts, USA). Sodium chloride (99%, S271-500, batch 225843) and sodium hydroxide (97%, S320-500, batch 241690) were purchased from Fisher Scientific Company (Pittsburgh, Pennsylvania, USA). Hydrochloric acid (HCl, 36.5 to 38.0 w/w %, A144-212, batch 235809) was purchased from Fisher Scientific Company (Pittsburgh, Pennsylvania, USA). All chemicals were used as purchased without further purification. 1X phosphate-buffered saline (PBS) was purchased from Gibco (Waltham, Massachusetts, USA). Water was purified with a Millipore Milli-Q Integral 10 water purification system (MilliporeSigma, Burlington, Massachusetts, USA).

### Preparation of amino acid solutions in neutral form

For amino acids soluble in their neutral form at 3 M, 3 mL of deionized water was added to a glass vial containing 15 mmol of the amino acid. Ultrasonic treatment was applied to facilitate dissolution, and the solution was then heated in a laboratory oven (Heratherm OMH60, Thermo Scientific, Langenselbold, Germany) at 80 ℃ to promote complete dissolution. Once fully dissolved, the solution was transferred to a 5 mL volumetric flask, and the final volume was adjusted precisely to 5 mL with deionized water. Refer to **Table 1** for the amino acids that are soluble in their neutral form at 3 M.

### Preparation of amino acid solutions in protonated form

2.5 mL of 12 M HCl was mixed with 3 mL of deionized water and added to a vial containing 30 mmol of the amino acid. Ultrasonic treatment was applied to facilitate the complete dissolution of the amino acid. Once fully dissolved, the mixture was transferred to a 10 mL volumetric flask, and the final volume was adjusted precisely to 10 mL with deionized water. Refer to **Table 1** for amino acids that are soluble in their protonated form at 3 M.

### Preparation of amino acid solutions in deprotonated form

A 10 M sodium hydroxide stock solution was prepared in a 100 mL volumetric flask. Next, 3 mL of 10 M NaOH solution was mixed with 3 mL of deionized water and added to a vial containing 30 mmol of the amino acid. Ultrasonic treatment was applied to facilitate the complete dissolution of the amino acid. Once fully dissolved, the mixture was transferred to a 10 mL volumetric flask, and the final volume was adjusted to 10 mL with deionized water. Refer to **Table 1** for amino acids that are soluble in their deprotonated form at 3 M.

### UV-Vis absorption and transmission spectroscopy

UV-Vis absorption spectra of amino acid solutions at a concentration of 0.5 mg/mL were measured using an Agilent Cary 6000i UV/Vis/NIR spectrophotometer (Agilent Technologies, Santa Clara, CA, USA). All amino acid samples except tryptophan were measured using a 1 mm optical path length quartz cuvette (high precision, two polished sides; Science Outlet, Weifang, China), while tryptophan was measured using a 10 μm optical path length quartz cuvette (Starna Cells Inc., Atascadero, CA, USA).

Furthermore, UV-Vis absorption spectra of arginine hydrochloride solutions at a series of low concentrations (50 mM, 100 mM, 250 mM, and 500 mM) were measured using an Agilent Cary 6000i UV/Vis/NIR spectrophotometer (Agilent Technologies, Santa Clara, CA, USA) equipped with a 10 μm optical path length quartz cuvette (Starna Cells Inc., Atascadero, CA, USA).

Additionally, transmission spectra of arginine hydrochloride solutions at higher concentrations (0.5 M, 1 M, 2 M, and 3 M) were acquired on the same Agilent Cary 6000i UV/Vis/NIR spectrophotometer using a 1 mm optical path length quartz cuvette (high precision, two polished sides; Science Outlet, Weifang, China).

### Refractometry measurements

Refractive index was measured using a digital refractometer (OPTi, Bellingham+ Stanley, Tunbridge Wells, UK). Prior to each measurement, the prism well was cleaned with deionized water and dried. A volume of 1 mL of the amino acid solution was applied to the prism well, and the refractive index was recorded automatically. Measurements were performed at a fixed wavelength (589 nm, sodium D-line equivalent) under ambient light conditions.

### Ellipsometry measurements

A Horiba Jobin Yvon UVISEL ellipsometer (Horiba Jobin Yvon, Longjumeau, France, 2015 Model) was used to measure the real and imaginary components of the complex refractive index of amino acid solutions. To prepare the sample, 320-grit sandpaper (SiC A-99, Partsmaster 881-5-0320, Dallas, Texas, USA) was attached to the bottom of the 25 × 20 × 5 mm cryomold (Sakura Finetek, Torrance, California, USA) using double-sided tape. Subsequently, 2.5 mL of the solution was added to the mold, forming a flat, reflective air-liquid interface. The real and imaginary refractive index spectra of the amino acid solution were measured in the wavelength range of 250 nm to 750 nm, with a reflection angle of 69.85°, a step size of 5 nm, and a dwell time of 200 ms.

### Osmolality and pH measurements

Osmolality was measured using a vapor pressure osmometer (Model 5600, ELITechGroup Inc., Logan, Utah, USA). Prior to the measurement, the instrument was calibrated using standard solutions according to the manufacturer’s instructions. The pH of the solutions was measured using a SevenCompact Duo pH/conductivity meter (Mettler Toledo, Columbus, Ohio, USA), which was calibrated with standard buffer solutions before each measurement. All measurements were performed at room temperature.

### Vertebrate animal subjects

All animal studies were conducted in compliance with procedures approved by the Stanford Institutional Animal Care and Use Committee (IACUC) and adhered to the guidelines outlined in the National Institutes of Health Guide for the Care and Use of Laboratory Animals. Male and female C57BL/6J mice were obtained from Jackson Labs (Sacramento, California, USA) and maintained at Stanford University’s Veterinary Service Center under controlled conditions (12 h light/dark cycle, 20-25 °C, 50-65% humidity). Animals were provided with standard bedding, nesting materials, food and water by the facility. Prior to experimentation, fur was removed using a depilatory cream (Nair, Body Cream, Church & Dwight, Ewing Township, New Jersey, USA). For *in vivo* clearing and imaging procedures, mice were anesthetized with 2.5% isoflurane and air as a carrying gas at a flow rate of 2 L/min. Animals were continuously monitored during procedures and throughout recovery in accordance with animal care protocols. Each experimental group included four mice (*n* = 4). At the study endpoint, the animals were humanely euthanized using carbon dioxide.

### Achieving optical transparency in dissected tissues

Mice (n = 4 each for the arginine hydrochloride treatment group and osmolarity-matched saline control group) were euthanized using carbon dioxide via inhalation, followed by cervical dislocation to ensure death. Nair (Body Cream, Church & Dwight, Ewing Township, New Jersey, USA) was applied generously to the abdominal area and allowed to remain for 5 minutes without rubbing before being wiped off with alcohol pads (Fisher Scientific Company, Fisher HealthCare, Pittsburgh, Pennsylvania, USA) to depilate the skin. A 10 mm × 20 mm rectangle of abdominal skin was excised using surgical scissors. The excised skin was placed in a Petri dish with the epidermis facing upward, and its corners were adhered to the bottom of the dish using Krazy Glue All Purpose Super Glue (Elmer’s Products, Atlanta, Georgia, USA). The Petri dish was positioned on a transparency film with a 1 × 1 mm grid pattern, which was placed on an LED light board inside a 37°C orbital shaker. The shaker remained stationary and was used solely for temperature control. The arginine hydrochloride solution or osmolarity-matched saline was carefully added to the Petri dish until it covered the excised mouse skin by at least 1 mm. The lid of the Petri dish was then closed and sealed with Parafilm. A color charge-coupled device (CCD) camera (DCU224C, Thorlabs) with an array size of 1280 × 1024 and a sensitivity range from 400 to 700 nm was set up. The camera was outfitted with a Xenon variable aperture f/0.95 lens (25 mm focal length). The camera was positioned to look down at the Petri dish, ensuring the plane of the grid pattern was in focus. The camera was configured with an exposure time of 14 ms. Images were acquired using the camera at 0 min, 30 min, 1 h, 3 h, and 10 h of soaking to document any changes in skin appearance over time.

### Achieving optical transparency in the abdomen of live mice

The arginine hydrochloride solution used for achieving in vivo optical transparency was prepared by dissolving 600 mg of arginine hydrochloride powder in 1 mL of deionized water. The mixture was incubated in an 80 °C oven and sonicated until complete dissolution was achieved. After cooling to room temperature, the final solution volume was approximately 1.4 mL, yielding a final concentration of 2.0 M.

Male and female C57BL/6J mice (4-5 weeks old, 11∼13 g, n = 4) from Jackson Labs (Sacramento, California, USA) were used for in vivo abdominal clearing and imaging experiments. Although mice older than 4 weeks or heavier than 15 g can also exhibit increased skin transparency following treatment with arginine hydrochloride, the accumulation of subcutaneous adipose tissue at these ages limits optical access, as the underlying fat remains opaque and obscures internal organs. Accordingly, for direct visualization of abdominal organs by the naked eye, younger mice (4-5 weeks, <15 g) are preferred.

Mice were anesthetized using isoflurane (Animal Anesthesia Vaporizer, RWD, Sugar Land, Texas, USA; Isoflurane, Dechra Veterinary Products, Overland Park, Kansas, USA) and maintained on a heating pad (Sunbeam, Boca Raton, Florida, USA). To maintain hydration, mice received a dorsal subcutaneous injection of saline (0.9% sodium chloride, USP; Hospira Inc., Lake Forest, Illinois, USA) at a dose of 20 μL per gram of body weight. Nair (Church & Dwight, Ewing Township, New Jersey, USA) was used for abdominal hair removal and abdominal skin exfoliation using a protocol modified from our previous report.^32^ Specifically, the depilatory cream was topically applied to the abdominal skin for 40 s with gentle massage to facilitate its penetration into the skin. The cream was then removed using dry Kimwipes tissues (Kimberly-Clark, Irving, Texas, USA), after which the skin was cleaned for 2 min using Kimwipe tissues soaked in water. Finally, the area was wiped with alcohol pads for three rounds to ensure complete removal of any residual cream. The entire procedure, from cream application to final cleaning, was completed within 5 min. For topical clearing, approximately 0.1 mL of arginine hydrochloride solution (2.0 M) was applied using a cotton-tipped applicator and massaged over an approximately 1 cm^2^ area for 10 min. During this period, additional solutions were applied using the same or a fresh applicator as needed to ensure continuous coverage of the clearing solution on the target area.

After sufficient transparency was achieved in the targeted abdominal region, excess solution was gently removed from the skin. The mouse was then imaged under white-light illumination for both photography and video recording. No glass slide or coverslip was required to flatten the skin surface, as evidenced by the images shown in **Fig. 6**. Notably, owing to the absence of coloration in arginine hydrochloride, internal organs such as the liver and intestines could be directly visualized through the transparent abdominal skin with the naked eye, without the need for specialized imaging equipment. Brightfield images were acquired using a home-built imaging system equipped with a color charge-coupled device (CCD) camera (DCU224C, Thorlabs, Newton, New Jersey, USA). The camera features an array size of 1280 × 1024 pixels and a spectral sensitivity range of 400-700 nm, with quantum efficiency exceeding 50% across this range. The system was fitted with a Schneider Xenon 25 mm f/0.95 lens for imaging the mouse abdomen. No optical filters were used. An exposure time of 50 ms was used for image acquisition, and videos were recorded at 15 frames per second in brightfield mode.

### Reversing optical transparency and chronic monitoring of live mice

After imaging, the mouse abdomen was gently rinsed with 5 mL of warm 1X PBS (Corning Inc., Corning, New York, USA) using a transfer pipette (13-711-9AMMD, FisherBrand, Pittsburgh, Pennsylvania, USA) to remove arginine hydrochloride absorbed in the skin and reverse the transparency effect. Delicate task wipers (Kimberly-Clark, Irving, Texas, USA) were used to prevent the surrounding fur from becoming wet, thereby minimizing the risk of hypothermia or infection. The abdomen was then cleaned with alcohol prep pads (22-363-750, FisherBrand, Pittsburgh, Pennsylvania, USA). A generous amount of Eucerin Advanced Repair Cream (Beiersdorf Inc., Stamford, Connecticut, USA) was applied to fully cover the treated area. The abdomen, with excess Eucerin, was then wrapped with surgical tape (3M Durapore, 1 inch × 30 feet, B014SPIK22, Saint Paul, Minnesota, USA) to protect the treated region. It is essential that the tape be wrapped such that the adhesive side adheres to itself; otherwise, the tape may detach once the mouse becomes active.

Following treatment, the mouse was returned to a cage with bedding placed partially over a heating pad (731-500-000R, Sunbeam, Boca Raton, Florida, USA), ensuring that only half of the cage was warmed so the mouse could move freely between heated and room-temperature areas. For longitudinal imaging of the abdominal skin on consecutive days after treatment, the surgical tape was removed immediately prior to imaging, after which Eucerin and surgical tape were reapplied. If imaging was not required every day, Eucerin and surgical tape were only replaced after the mouse had damaged the surgical tape.

### Animal preparation for tissue collection and histopathology

Age-matched C57BL/6J (male and female, 6 weeks, 17-23 g) mice were used for all experiments. 4 mice (2 male and 2 female mice) were depilated, exfoliated, treated with arginine hydrochloride (2.0 M) in the abdominal skin, and subsequently reversed for the transparency effect as described in the “Achieving optical transparency in the abdomen of live mice” section. The mice were monitored and given the same recovery treatment with Eucerin as described above. The mice were then submitted to the Stanford Veterinary Service Center Necropsy Laboratory for histology, and to the Animal Diagnostic Laboratory for blood chemistry and hematology assays after 24 h of recovery, as detailed below in “Tissue collection and histopathology”. The same exact process was done on another 2 male and 2 female mice using phosphate buffered saline (PBS) as a control. The same process was repeated on another 8 mice (4 male and 4 female, 17∼29 g evenly divided between the arginine hydrochloride group and the PBS control group, with an equal number of animals of each sex in each group), except these mice were submitted to the Veterinary Service Center after a 14-day recovery period.

### Tissue collection and histopathology

Mice were euthanized by CO_2_ asphyxiation and cardiac exsanguination. Terminal cardiac blood was collected for complete blood counts (CBC) and serum chemistry evaluation through the Stanford Animal Diagnostic Laboratory. The abdominal skin, subcutaneous tissue, and whole-body (heart, liver, gallbladder, spleen, kidney, adrenal gland, pancreas, salivary gland, thymus, lung, esophagus, trachea, thyroid, tongue, reproductive tract, urinary bladder, brain, and gastrointestinal tract) were collected and immersion fixed in 10% neutral buffered formalin for 72 h. Following fixation, two longitudinal sections of abdominal skin were collected along the ventral midline, parallel to hair follicle growth. Formalin-fixed samples were submitted to HistoTec Laboratories (Hayward, CA), processed routinely, embedded in paraffin, sectioned, and stained with hematoxylin and eosin (H&E). H&E samples were assessed by a board-certified veterinary pathologist, evaluating whole-body organ systems (listed above) for systemic toxicity and scoring of the abdominal skin with subcutaneous tissues. The epidermis/dermis was evaluated using an ordinal scale (scores 0-4) where scores reflect the percentage of tissue exhibiting a given finding. Histologic features accounted for are mucosal ulceration, follicular and dermal necrosis +/- inflammation, acanthosis and hyperkeratosis, serocellular crusts, dermal fibrosis, follicular and adnexal dropout or disorganization, and lymphoplasmacytic dermatitis. Scores were assigned as follows: 1: ≤ 25% of evaluated section, 2: 25-49% of evaluated section, 3: 50-74% of evaluated section, 4: ≥ 75% of evaluated section. The panniculus and deep muscle were evaluated using a binary scale (scores 0-1) denoting the presence or absence of inflammation: 1 = yes and 0 = no. Scores for each layer were totaled, resulting in a final score of 0-6.

### Statistics and reproducibility

All experiments were independently repeated with the exact number of repeats provided below and in the relevant sections of the procedure. Comparison between groups was evaluated using ordinary one-way ANOVA, with *P* < 0.05 considered statistically significant. The sample size for each experiment was determined by power analysis to ensure statistical rigour for all comparisons:^54^

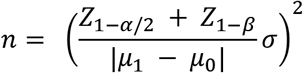

where *α* = 0.05, *Z*_1−*α*/2_ = 1.96, *Z*_1−*β*_ = 0.84, *β* = 0.20, *μ*_1_ is the mean of the outcome variable for the experimental condition, *μ*_0_ is the mean of the outcome variable for the control and *σ* is the s.d. of the outcome variable.

Achieving optical transparency in dissected *ex vivo* abdominal skin was independently replicated in 4 mice per group for both the arginine hydrochloride treatment group and the osmolarity-matched saline control group, resulting in a total of 8 mice. Achieving and reversing optical transparency in the abdominal skin of live mice was independently replicated in 4 mice. Tissue collection and histopathological analyses were performed in 4 mice per experimental condition, resulting in a total of 16 mice. Data were assumed to be normally distributed, although this was not formally tested. No data were excluded from the analyses. Experimental animals were healthy and in a comparable physiological state before the experiments and were randomly assigned to experimental groups. Investigators were blinded to group allocation during sample collection and data analysis.

## Data availability

The data that support the findings of this study are available within this Article and its Supplementary Information. Source data are provided with this paper.

## Acknowledgments

G.H. acknowledges two awards from the NIH (grant nos. 1R34NS127103 and R01NS126076), an NSF CAREER award (grant no. 2045120), an NSF EAGER award (grant no. 2217582), a Rita Allen Foundation Scholars Award, a Firmenich Next Generation Faculty Scholarship from the Firmenich Foundation, two gifts from the Pinetops Foundation, a teacher-scholar award from the Camille and Henry Dreyfus Foundation, a Wu Tsai Synthetic Neurobiology Grant, the Big Ideas in Neuroscience grant, the Alfred P. Sloan Research Fellowship, and the Pershing Square Foundation MIND Prize. L.-Y.Z. acknowledges a Knight-Hennessy Fellowship. H.C. acknowledges the support of a Stanford Interdisciplinary Graduate Fellowship as a David L. Sze and Kathleen Donohue Interdisciplinary Fellow. V.C. acknowledges the Stanford Graduate Fellowship. C.H.C.K. acknowledges the National Science Foundation Graduate Research Fellowships program (grant no. 1656518) and the Wu Tsai Neuroscience NeuroTech Training program. Part of this work was performed at nano@stanford RRID:SCR_026695. Some schematics were created with BioRender.com.

## Contributions

S.Z., Z.L., L.-Y.Z. and G.H. conceived and designed the project. S.Z., Z.L., L.-Y.Z., X.H., H.C., D.M. and T.M.Y. prepared amino acid solutions and performed their comprehensive optical characterizations. S.Z. and X.H. performed pH and osmolality measurements of amino acid solutions. L.-Y.Z. performed ex vivo skin clearing experiments. S.Z., V.C. and C.H.C.K. performed in vivo mouse experiments for abdominal clearing, reversal, and monitoring. S.Z. and A.B. prepared animals for tissue collection and histopathology. K.M.C. and P.M.W. performed complete blood counts, serum chemistry evaluation, and histopathology for tissue samples. S.Z., Z.L., L.-Y.Z., A.B. and G.H. analysed the data. All authors contributed to the writing of the paper.

## Competing interests

The authors declare no competing interests.

## Notes

### Competing Interest Statement

The authors have declared no competing interest.

## References

(1) Ntziachristos, V. Going Deeper than Microscopy: The Optical Imaging Frontier in Biology. Nat. Methods 2010, 7 (8), 603–614.

(2) Fenno, L.; Yizhar, O.; Deisseroth, K. The Development and Application of Optogenetics. Annu. Rev. Neurosci. 2011, 34, 389–412.

(3) Sutton, P. A.; van Dam, M. A.; Cahill, R. A.; Mieog, S.; Polom, K.; Vahrmeijer, A. L.; van der Vorst, J. Fluorescence-Guided Surgery: Comprehensive Review. BJS Open 2023, 7 (3). 10.1093/bjsopen/zrad049.

(4) Li, X.; Lovell, J. F.; Yoon, J.; Chen, X. Clinical Development and Potential of Photothermal and Photodynamic Therapies for Cancer. Nat. Rev. Clin. Oncol. 2020, 17 (11), 657–674.

(5) Liu, X.; Wang, W.; Artman, B.; Diao, J.; Zhao, Y.; He, W.; Yu, S.; Tang, K. W. K.; Yao, M.; Gu, C.; Song, B.; Wang, H. Multicolored, Sonosensitizer-Optimized Organic Mechanoluminescent Nanoparticles for Functional Sono-Optogenetics. J. Am. Chem. Soc. 2026, 148 (16), 16809–16820.

(6) Wang, W.; Kevin Tang, K. W.; Pyatnitskiy, I.; Liu, X.; Shi, X.; Huo, D.; Jeong, J.; Wynn, T.; Sangani, A.; Baker, A.; Hsieh, J.-C.; Lozano, A. R.; Artman, B.; Fenno, L.; Buch, V. P.; Wang, H. Ultrasound-Induced Cascade Amplification in a Mechanoluminescent Nanotransducer for Enhanced Sono-Optogenetic Deep Brain Stimulation. ACS Nano 2023, 17 (24), 24936–24946.

(7) Li, P.; Zhang, J.; Hayashi, H.; Yue, J.; Li, W.; Yang, C.; Sun, C.; Shi, J.; Huberman-Shlaes, J.; Hibino, N.; Tian, B. Monolithic Silicon for High Spatiotemporal Translational Photostimulation. Nature 2024, 626 (8001), 990–998.

(8) Jiang, Y.; Li, X.; Liu, B.; Yi, J.; Fang, Y.; Shi, F.; Gao, X.; Sudzilovsky, E.; Parameswaran, R.; Koehler, K.; Nair, V.; Yue, J.; Guo, K.; Fang, Y.; Tsai, H.-M.; Freyermuth, G.; Wong, R. C. S.; Kao, C.-M.; Chen, C.-T.; Nicholls, A. W.; Wu, X.; Shepherd, G. M. G.; Tian, B. Rational Design of Silicon Structures for Optically Controlled Multiscale Biointerfaces. Nat Biomed Eng 2018, 2 (7), 508–521.

(9) Jacques, S. L. Optical Properties of Biological Tissues: A Review. Phys. Med. Biol. 2013, 58 (11), R37–R61.

(10) Tuchin, V. V. Tissue Optics and Photonics: Light-Tissue Interaction. Journal of Biomedical Photonics & Engineering. 2015, pp 98–134. 10.18287/jbpe-2015-1-2-98.

(11) Ueda, H. R.; Ertürk, A.; Chung, K.; Gradinaru, V.; Chédotal, A.; Tomancak, P.; Keller, P. J. Tissue Clearing and Its Applications in Neuroscience. Nat. Rev. Neurosci. 2020, 21 (2), 61–79.

(12) Rakhilin, N.; Garrett, A.; Eom, C.-Y.; Chavez, K. R.; Small, D. M.; Daniel, A. R.; Kaelberer, M. M.; Mejooli, M. A.; Huang, Q.; Ding, S.; Kirsch, D. G.; Bohórquez, D. V.; Nishimura, N.; Barth, B. B.; Shen, X. An Intravital Window to Image the Colon in Real Time. Nat. Commun. 2019, 10 (1), 5647.

(13) Sahasrabudhe, A.; Rupprecht, L. E.; Orguc, S.; Khudiyev, T.; Tanaka, T.; Sands, J.; Zhu, W.; Tabet, A.; Manthey, M.; Allen, H.; Loke, G.; Antonini, M.-J.; Rosenfeld, D.; Park, J.; Garwood, I. C.; Yan, W.; Niroui, F.; Fink, Y.; Chandrakasan, A.; Bohórquez, D. V.; Anikeeva, P. Multifunctional Microelectronic Fibers Enable Wireless Modulation of Gut and Brain Neural Circuits. Nat. Biotechnol. 2024, 42 (6), 892–904.

(14) Zon, L. I.; Peterson, R. T. In Vivo Drug Discovery in the Zebrafish. Nat. Rev. Drug Discov. 2005, 4 (1), 35–44.

(15) Taboada, C.; Delia, J.; Chen, M.; Ma, C.; Peng, X.; Zhu, X.; Jiang, L.; Vu, T.; Zhou, Q.; Yao, J.; O’Connell, L.; Johnsen, S. Glassfrogs Conceal Blood in Their Liver to Maintain Transparency. Science 2022, 378 (6626), 1315–1320.

(16) Inyushin, M.; Meshalkina, D.; Zueva, L.; Zayas-Santiago, A. Tissue Transparency in Vivo. Molecules 2019, 24 (13), 2388.

(17) Benedek, G. B. Theory of Transparency of the Eye. Appl. Opt. 1971, 10 (3), 459–473.

(18) Zhao, H.; Brown, P. H.; Schuck, P. On the Distribution of Protein Refractive Index Increments. Biophys. J. 2011, 100 (9), 2309–2317.

(19) McMEEKIN, T. L.; Groves, M. L.; Hipp, N. J. Refractive Indices of Amino Acids, Proteins, and Related Substances. In *Advances in Chemistry*; Advances in Chemistry Series; AMERICAN CHEMICAL SOCIETY: WASHINGTON, D.C., 1964; pp 54–66.

(20) McMeekin, T. L.; Wilensky, M.; Groves, M. L. Refractive Indices of Proteins in Relation to Amino Acid Composition and Specific Volume. Biochem. Biophys. Res. Commun. 1962, 7 (2), 151–156.

(21) Kappé, G.; Purkiss, A. G.; van Genesen, S. T.; Slingsby, C.; Lubsen, N. H. Explosive Expansion of Betagamma-Crystallin Genes in the Ancestral Vertebrate. J. Mol. Evol. 2010, 71 (3), 219–230.

(22) Kiteto, M. K.; Mecha, C. A. Insight into the Bouguer-Beer-Lambert Law: A Review. Sustainable Chemical Engineering 2024, 567–587.

(23) Mayerhöfer, T. G.; Pipa, A. V.; Popp, J. Beer’s Law-Why Integrated Absorbance Depends Linearly on Concentration. Chemphyschem 2019, 20 (21), 2748–2753.

(24) Nagle, J. K. Atomic Polarizability and Electronegativity. J. Am. Chem. Soc. 1990, 112 (12), 4741–4747.

(25) Gund, P. Guanidine, Trimethylenemethane, and “Y-Delocalization.” Can Acyclic Compounds Have “Aromatic” Stability? J. Chem. Educ. 1972, 49 (2), 100.

(26) Zhao, H.; Brown, P. H.; Magone, M. T.; Schuck, P. The Molecular Refractive Function of Lens γ-Crystallins. J. Mol. Biol. 2011, 411 (3), 680–699.

(27) Kühnel, K.; Ke, N.; Cryle, M. J.; Sligar, S. G.; Schuler, M. A.; Schlichting, I. Crystal Structures of Substrate-Free and Retinoic Acid-Bound Cyanobacterial Cytochrome P450 CYP120A1. Biochemistry 2008, 47 (25), 6552–6559.

(28) Uhlhorn, S. R.; Borja, D.; Manns, F.; Parel, J.-M. Refractive Index Measurement of the Isolated Crystalline Lens Using Optical Coherence Tomography. Vision Res. 2008, 48 (27), 2732–2738.

(29) Couto Oliveira, L. M.; Tuchin, V. V. The Optical Clearing Method: A New Tool for Clinical Practice and Biomedical Engineering, 1st ed.; SpringerBriefs in physics; Springer Nature: Cham, Switzerland, 2019.

(30) Inagaki, S.; Nakagawa-Tamagawa, N.; Huynh, N. Z.; Kambe, Y.; Yagasaki, R.; Manita, S.; Fujimoto, S.; Noda, T.; Mori, M.; Teranishi, A.; Takeshima, H.; Ishikawa, K.; Naitou, Y.; Yokoyama, T.; Sakamoto, M.; Hayashi, K.; Kitamura, K.; Tagawa, Y.; Okuda, S.; Sato, T. K.; Imai, T. Isotonic and Minimally Invasive Optical Clearing Media for Live Cell Imaging Ex Vivo and in Vivo. Nat. Methods 2026. 10.1038/s41592-026-03023-y.

(31) Hou, X.; Cai, S.; Cui, H.; Liu, Z.; Zhao, S.; Zhang, L.-Y.; Baghdasaryan, A.; Crunkleton, V.; Brongersma, M. L.; Hong, G. Tartrazine Clears Live Cells While Preserving Viability at High Refractive Indices and Osmolality. Bioconjug. Chem. 2026, No. acs.bioconjchem.6c00164. 10.1021/acs.bioconjchem.6c00164.

(32) Keck, C. H. C.; Schmidt, E. L.; Zhao, S.; Liu, Z.; Zhang, L.-Y.; Cui, M.; Chen, X.; Wang, C.; Cui, H.; Brongersma, M. L.; Hong, G. Achieving Transient and Reversible Optical Transparency in Live Mice with Tartrazine. Nat. Protoc. 2025. 10.1038/s41596-025-01187-z.

(33) Ou, Z.; Duh, Y.-S.; Rommelfanger, N. J.; Keck, C. H. C.; Jiang, S.; Brinson, K., Jr; Zhao, S.; Schmidt, E. L.; Wu, X.; Yang, F.; Cai, B.; Cui, H.; Qi, W.; Wu, S.; Tantry, A.; Roth, R.; Ding, J.; Chen, X.; Kaltschmidt, J. A.; Brongersma, M. L.; Hong, G. Achieving Optical Transparency in Live Animals with Absorbing Molecules. Science 2024, 385 (6713), eadm6869.

(34) Keck, C. H. C.; Schmidt, E. L.; Roth, R. H.; Floyd, B. M.; Tsai, A. P.; Garcia, H. B.; Cui, M.; Chen, X.; Wang, C.; Park, A.; Zhao, S.; Liao, P. A.; Casey, K. M.; Reineking, W.; Cai, S.; Zhang, L.-Y.; Yang, Q.; Yuan, L.; Baghdasaryan, A.; Lopez, E. R.; Cooper, L.; Cui, H.; Esquivel, D.; Brinson, K.; Chen, X.; Wyss-Coray, T.; Coleman, T. P.; Brongersma, M. L.; Bertozzi, C. R.; Wang, G. X.; Ding, J. B.; Hong, G. Color-Neutral and Reversible Tissue Transparency Enables Longitudinal Deep-Tissue Imaging in Live Mice. Proc. Natl. Acad. Sci. U. S. A. 2025, 122 (35), e2504264122.

(35) Miller, D. A.; Xu, Y.; Highland, R.; Nguyen, V. T.; Brown, W. J.; Hong, G.; Yao, J.; Wax, A. Enhanced Penetration Depth in Optical Coherence Tomography and Photoacoustic Microscopy in Vivo Enabled by Absorbing Dye Molecules. Optica 2025, 12 (1), 24–30.

(36) Seong, D.; Yun, S.; Han, S.; Biswas, S.; Kim, B.; Remlova, E.; Razansky, D.; Kim, J.; Ou, Z.; Jeon, M. Beyond the Skin Barrier: Optical Clearing Enables Non-Invasive Cortex-Wide Optical Coherence Angiography in Mice in-Vivo. bioRxiv, 2026. 10.64898/2026.03.02.709062.

(37) Crunkleton, V.; Hong, G. In Vivo Tissue Clearing with Tartrazine and Other Dye Molecules. Commun. Biol. 2026, 9 (1). 10.1038/s42003-026-10610-4.

(38) Callaway, J. P.; Bharadwaj, S.; Li, Z.; Trenkle, A. S.; Kwong, G. A.; Hong, G.; Robles, F. E. Tartrazine-Enabled Enhanced Image Depth Penetration with Quantitative Oblique Back-Illumination Microscopy. Opt. Lett. 2026, 51 (15), 4072–4075.

(39) Zuo, T.; Tao, C.; Liu, X. Absorbing Molecules as Optical Clearing Agents Improve the Resolution and Sensitivity of Photoacoustic Microscopy. Opt. Lett. 2025, 50 (7), 2282–2285.

(40) Jia, C.; Zhang, Z.; Shen, Y.; Hou, W.; Zhao, J.; Luo, J.; Chen, H.; Qi, D.; Yao, Y.; Deng, L.; Ma, H.; Sun, Z.; Zhang, S. Tartrazine-Enabled Optical Clearing for in Vivo Optical Resolution Photoacoustic Microscopy. Biomed. Opt. Express 2025, 16 (6), 2504–2515.

(41) Narawane, A.; Trout, R.; Viehland, C.; Kuo, A. N.; Vajzovic, L.; Dhalla, A.-H.; Toth, C. A. Optical Clearing with Tartrazine Enables Deep Transscleral Imaging with Optical Coherence Tomography. J. Biomed. Opt. 2024, 29 (12), 120501.

(42) Kim, S.-J.; Jo, K.; Park, K.; Kim, J. H.; Cho, J. H.; Choi, I.; Hahn, S. K. Strain-programmable Luminescent Adhesive Patch with Tartrazine-mediated Optical Skin Clearing for Photochemical Tissue Bonding. Adv. Funct. Mater. 2026, No. e29087. 10.1002/adfm.202529087.

(43) Surkov, Y.; Timoshina, P.; Uvakin, I.; Shushunova, N.; Konovalov, A.; Kozlov, I.; Piavchenko, G.; Telyshev, D.; Meglinski, I.; Kuznetsov, S.; Tuchin, V. Computer-Guided Optical Clearing for Transcranial Laser Speckle Imaging of Cortical Blood Flow through Synergistic Tartrazine-Induced Cranial Bone Transparency. J. Innov. Opt. Health Sci. 2025, No. 2540002. 10.1142/s1793545825400024.

(44) Yuan, N.; Ragab, S.; Chavez, L.; Pandey, V.; Intes, X. Evaluating Tartrazine as an Optical Clearing Agent for Fluorescence Lifetime Imaging. Opt. Lett. 2025, 50 (24), 7588–7591.

(45) Sun, T.; Su, J.; Zhao, Y.; Tie, X. Enhancing the Efficiency of Achieving Optical Transparency in Live Animals Using Absorbing Molecules. J. Biomed. Opt. 2026, 31 (5), 054702.

(46) Asante-Asare, D.; Kajla, R.; Gangaraju, M.; Taboada, E.; Argueta, A.; Gora, A.; Sehgal, G. K.; Nguyen, M.; Shabbir, M. W.; Slinker, J. D.; Wang, K. K.-H.; Ou, Z. Kinetics of Transient Tissue Transparency via Refractive Index Engineering (T3RIE) for in Vivo Optical Imaging. Chem. Eur. J. 2026, No. e03667, e03667.

(47) Breglia, R. J.; Ward, C. O.; Jarowski, C. I. Effect of Selected Amino Acids on Ethanol Toxicity in Rats. J. Pharm. Sci. 1973, 62 (1), 49–55.

(48) Wu, G.; Meininger, C. J.; McNeal, C. J.; Bazer, F. W.; Rhoads, J. M. Role of L-Arginine in Nitric Oxide Synthesis and Health in Humans. Adv. Exp. Med. Biol. 2021, 1332, 167–187.

(49) Myung, D.; Zhao, S.; Manning, E. C. H.; Brongersma, M.; Winetraub, Y.; Sarin, K.; Hong, G. Achieving Optical Transparency in Human Skin with Absorbing Molecules. In Dynamics and Fluctuations in Biomedical Photonics XXIII; Tuchin, V. V., Leahy, M. J., Wang, R. K., Eds.; SPIE, 2026; p 36.

(50) Chakrabarti, A. C. Permeability of Membranes to Amino Acids and Modified Amino Acids: Mechanisms Involved in Translocation. Amino Acids 1994, 6 (3), 213–229.

(51) Mitchell, D. J.; Kim, D. T.; Steinman, L.; Fathman, C. G.; Rothbard, J. B. Polyarginine Enters Cells More Efficiently than Other Polycationic Homopolymers. J. Pept. Res. 2000, 56 (5), 318–325.

(52) Wender, P. A.; Mitchell, D. J.; Pattabiraman, K.; Pelkey, E. T.; Steinman, L.; Rothbard, J. B. The Design, Synthesis, and Evaluation of Molecules That Enable or Enhance Cellular Uptake: Peptoid Molecular Transporters. Proc. Natl. Acad. Sci. U. S. A. 2000, 97 (24), 13003–13008.

(53) MacCallum, J. L.; Bennett, W. F. D.; Tieleman, D. P. Transfer of Arginine into Lipid Bilayers Is Nonadditive. Biophys. J. 2011, 101 (1), 110–117.

(54) Cohen, J. Statistical Power Analysis for the Behavioral Sciences; Academic Press, 2013.

(55) Prasad, S.; Mandal, I.; Singh, S.; Paul, A.; Mandal, B.; Venkatramani, R.; Swaminathan, R. Near UV-Visible Electronic Absorption Originating from Charged Amino Acids in a Monomeric Protein. Chem. Sci. 2017, 8 (8), 5416–5433.

