## Supplementary Information for "Refractive index modulation by ultraviolet absorption of canonical amino acids for *in vivo* optical transparency"

### Table of Contents

Supplementary Movie Legends……………………………………………………………….2

Supplementary Figures S1-S15…………………………………………….………………...3

Supplementary Table S1…………………………………….…………………………..………20

Supplementary References…………………………………………………………………...22

**Supplementary Movie Legends**

**Movie S1.** Representative movie of a live mouse with abdominal wall transparency induced by topical application of arginine hydrochloride. The mouse shown in this movie is the same as in **Fig. 6b**. The movie is shown at 1× speed.

**Movie S2.** Representative movie of another live mouse with abdominal wall transparency induced by topical application of arginine hydrochloride. The mouse shown in this movie is the same as in **Fig. 6c**. The movie is shown at 1× speed.

### Supplementary Figures

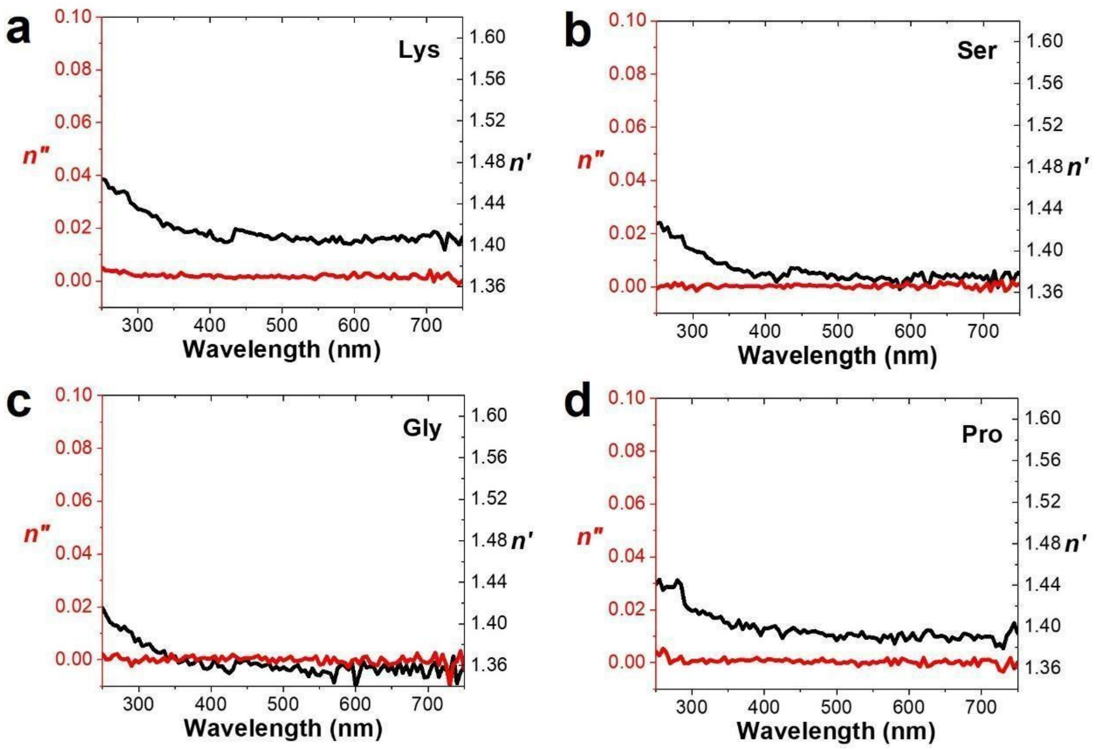


**Fig. S1.** Spectroscopic ellipsometry of (**a**) lysine, (**b**) serine, (**c**) glycine, and (**d**) proline in their original form. Spectra were measured at 3 M for all solutions. All other amino acids cannot be dissolved in water in their original form at 3 M.

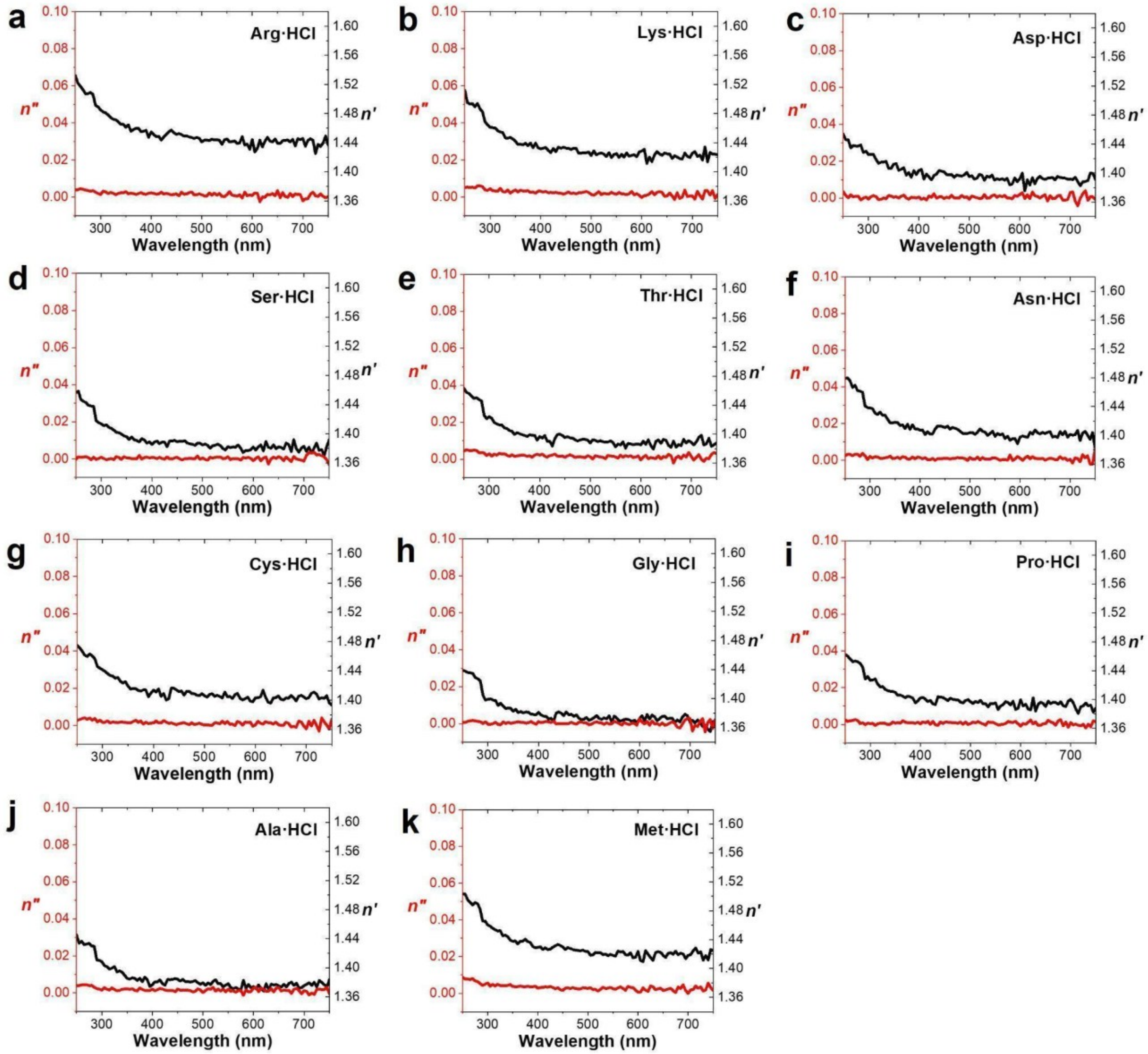


**Fig. S2.** Spectroscopic ellipsometry of (**a**) arginine, (**b**) lysine, (**c**) aspartic acid, (**d**) serine, (**e**) threonine, (**f**) asparagine, (**g**) cysteine, (**h**) glycine, (**i**) proline, (**j**) alanine, and (**k**) methionine in their protonated forms. Spectra were measured at 3 M for all solutions. All other amino acids cannot be dissolved in water in their protonated form at 3 M.

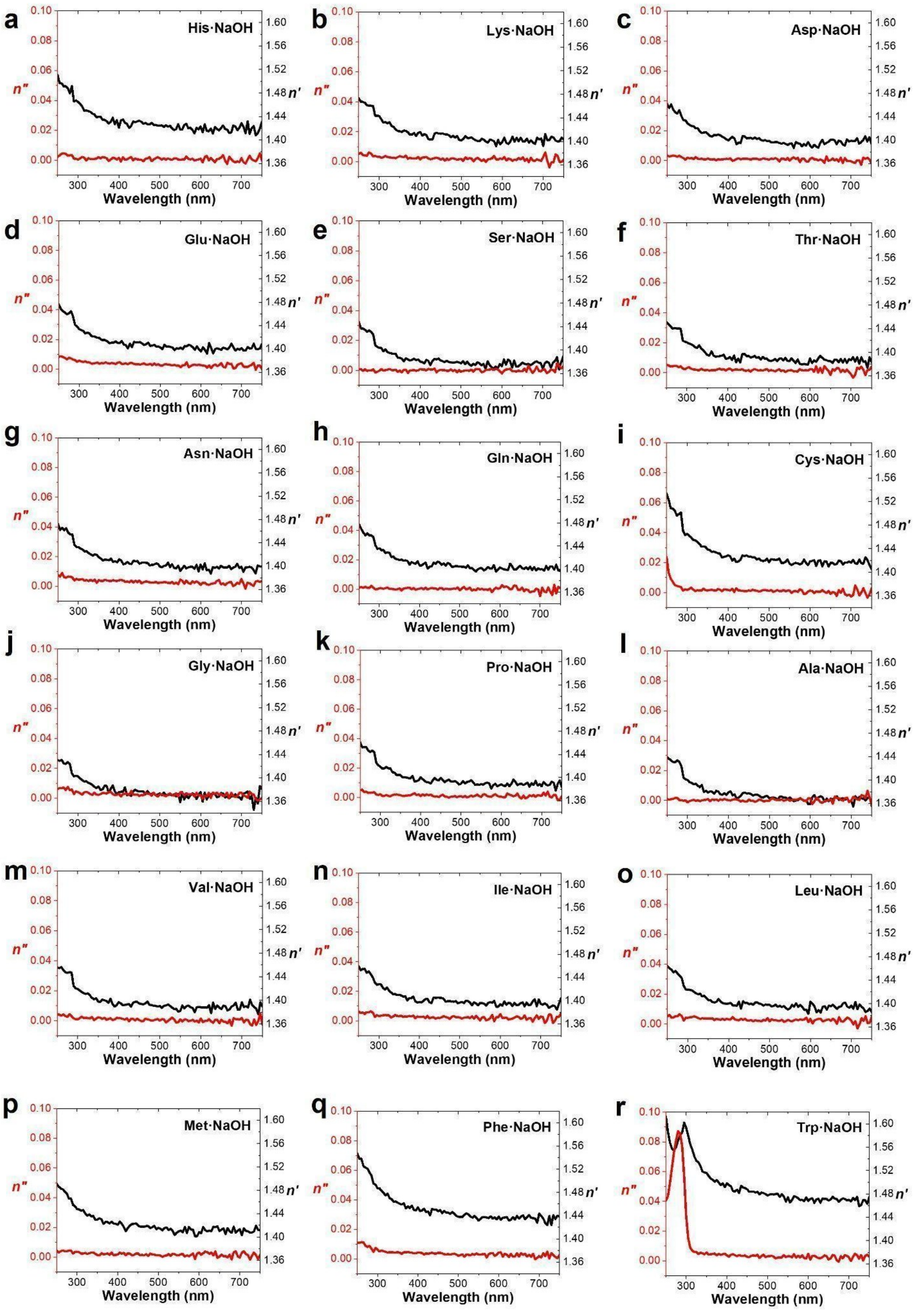
a
His·NaOH
b
Lys·NaOH
c
Asp·NaOH
d
Glu·NaOH
e
Ser·NaOH
f
Thr·NaOH
g
Asn·NaOH
h
Gln·NaOH
i
Cys·NaOH
j
Gly·NaOH
k
Pro·NaOH
l
Ala·NaOH
m
Val·NaOH
n
Ile·NaOH
o
Leu·NaOH
p
Met·NaOH
q
Phe·NaOH
r
Trp·NaOH
n″
n′
Wavelength (nm)
0.00 0.02 0.04 0.06 0.08 0.10
1.36 1.40 1.44 1.48 1.52 1.56 1.60
300 400 500 600 700

**Fig. S3.** Spectroscopic ellipsometry of (**a**) histidine, (**b**) lysine, (**c**) aspartic acid, (**d**) glutamic acid, (**e**) serine, (**f**) threonine, (**g**) asparagine, (**h**) glutamine, (**i**) cysteine, (**j**) glycine, (**k**) proline, (**l**) alanine, (**m**) valine, (**n**) isoleucine, (**o**) leucine, (**p**) methionine, (**q**) phenylalanine, and (**r**) tryptophan in their deprotonated forms. Spectra were measured at 3 M for all solutions. All other amino acids cannot be dissolved in water in their deprotonated form at 3 M.

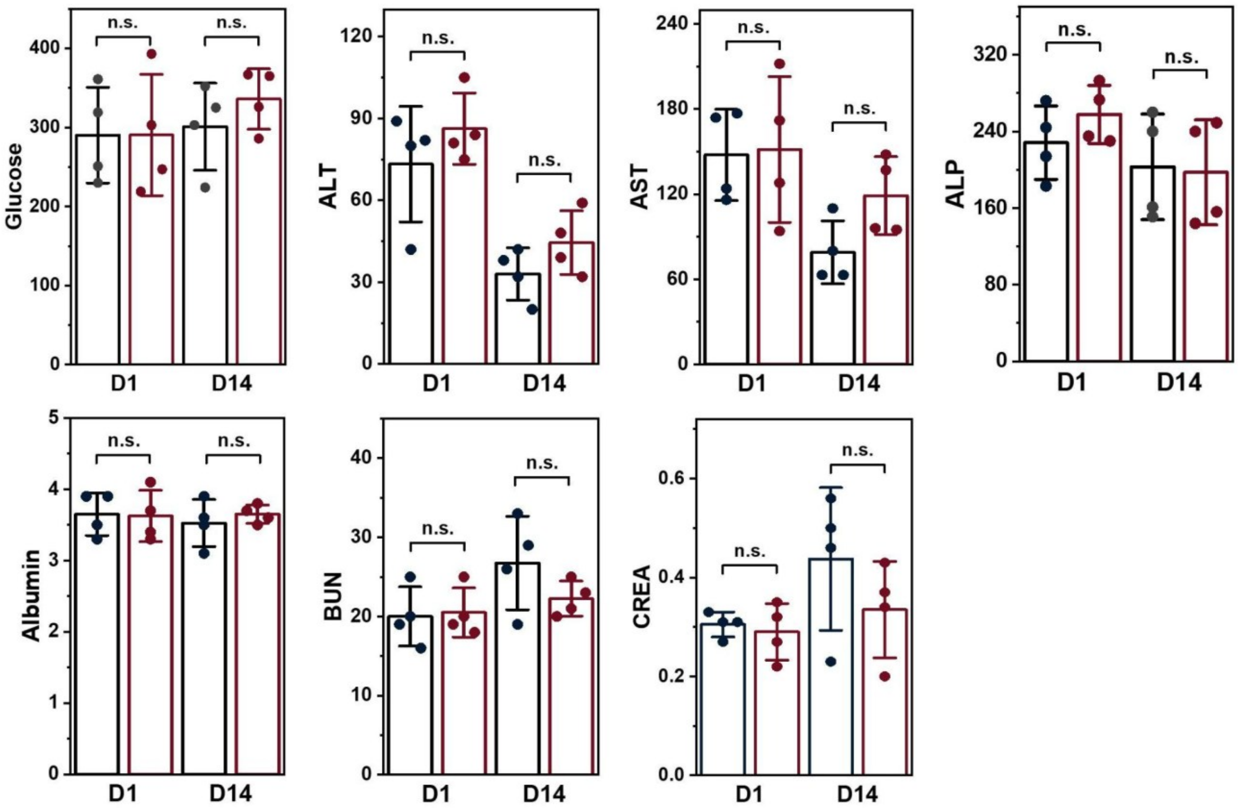


**Fig. S4.** Blood chemistry test results of experimental mouse groups at 24 h (D1) and 2 weeks (D14) following topical application of 2 M Arg·HCl solution to the abdominal area (n = 4 per time point, red). Mice treated with saline alone were used as the control group (n = 4 per time point, black). The test results of glucose (mg/dL), alanine aminotransferase (ALT; U/L), aspartate aminotransferase (AST; U/L), alkaline phosphatase (ALP; IU/L), albumin (g/dL), blood urea nitrogen (BUN; mg/dL) and creatinine (CREA; mg/dL) represent standard deviation (SD) of four repeated experiments. Bar graphs are presented as mean values ± SD. The data were analyzed by Tukey's test.

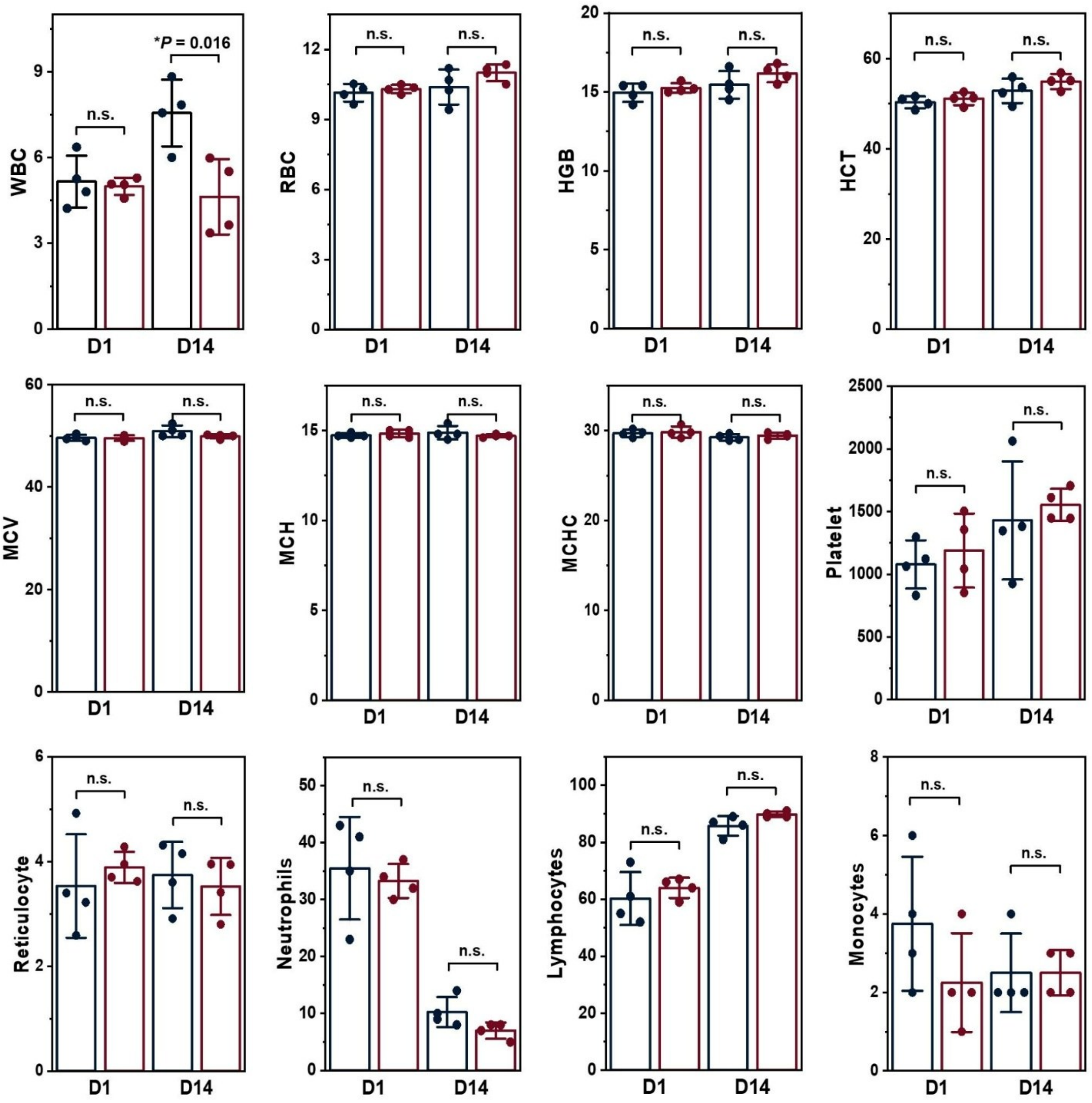


**Fig. S5.** Complete blood count (CBC) results of experimental mouse groups at 24 h (D1) and 2 weeks (D14) following topical application of 2 M Arg·HCl solution to the abdominal area (n = 4 per time point, red). Mice treated with saline alone were used as the control group (n = 4 per time point, black). The test results of white blood cells (WBC; K/μL), red blood cells (RBC; M/μL), hemoglobin (HGB; gm/dL), hematocrit (HCT; %), mean corpuscular volume (MCV; fL), mean corpuscular hemoglobin (MCH; pg), mean corpuscular hemoglobin concentration (MCHC; g/dL), platelet (K/μL), reticulocyte (%), neutrophils (%), lymphocytes (%), and monocytes (%) represent standard deviation (SD) of four repeated experiments. Bar graphs are presented as mean values ± SD. Note that for the CBC differential (i.e., neutrophils, lymphocytes, and monocytes), a manual WBC differential was performed for each sample. Specifically, a peripheral blood smear was reviewed, and 100 WBCs were counted manually. Consequently, the differential percentages for these cell types are reported as whole numbers (i.e., in 1% increments). The data were analyzed by Tukey's test. Absolute P value was derived using One-Way ANOVA.

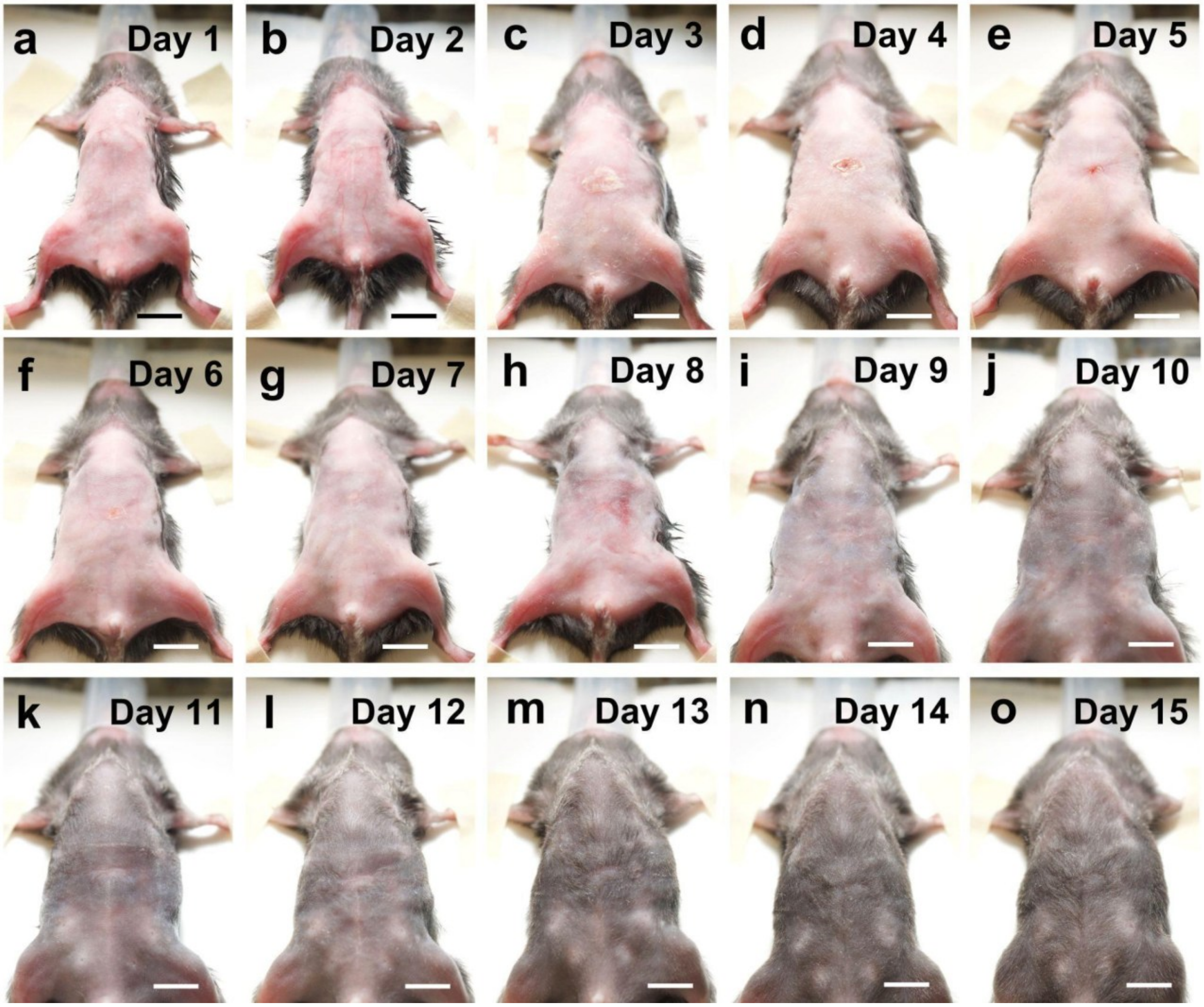


**Fig. S6.** Photographs showing the process of skin recovery in 15 days after achieving and reversing optical transparency induced by arginine hydrochloride in the abdomen of live mice. The same animal depicted in **Fig. 6c** is presented here. Scale bars: 1 cm.

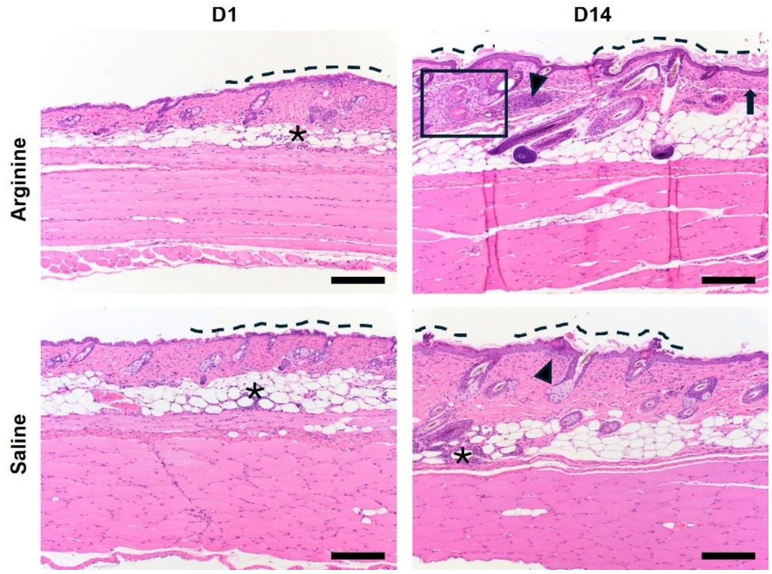


**Fig. S7.** Histology images of H&E-stained abdominal skin tissue sections from experimental mouse groups at 24 h (D1) and 2 weeks (D14) following topical application of 2 M Arg·HCl solution to the abdominal area and reversing the transparency effect with saline (n = 4 per time point). Mice treated with saline alone were used as the control group (n = 4 per time point). For 24 h (D1) images, the dashed line indicates ulceration with serocellular crusts. For 2 week (D14) images, the dashed line indicates acanthosis with hyperkeratosis, arrowheads indicate inflammation within the dermis, the arrow indicates dermal fibrosis, and the box indicates follicular and adnexal disorganization. For both time points, asterisks indicate inflammation within the panniculus. The presence of these minor skin changes in both the arginine-treated and saline-treated groups suggests that they result from mechanical rubbing rather than from the clearing agent, arginine hydrochloride, itself. Scale bars are 200 μm.

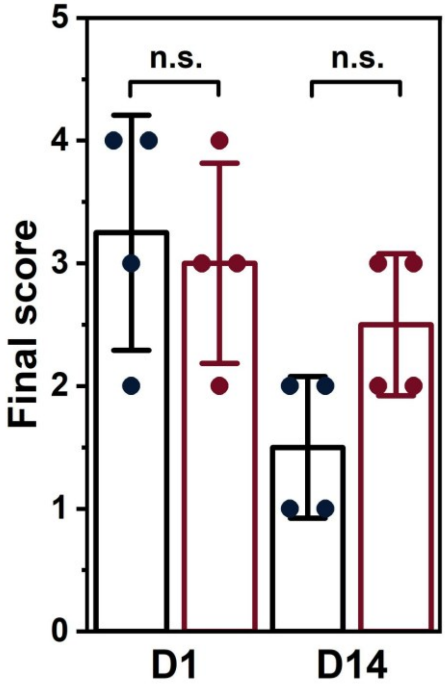


**Fig. S8.** Skin histology scoring. The final histological score represents the sum of the scores assigned to epidermal/dermal changes, panniculitis, and myositis. Black bars represent the control group, while red bars represent the experimental group. Data represent standard deviation (SD) of four repeated experiments. Bar graphs are presented as mean values ± SD. The data were analyzed by Tukey's test.

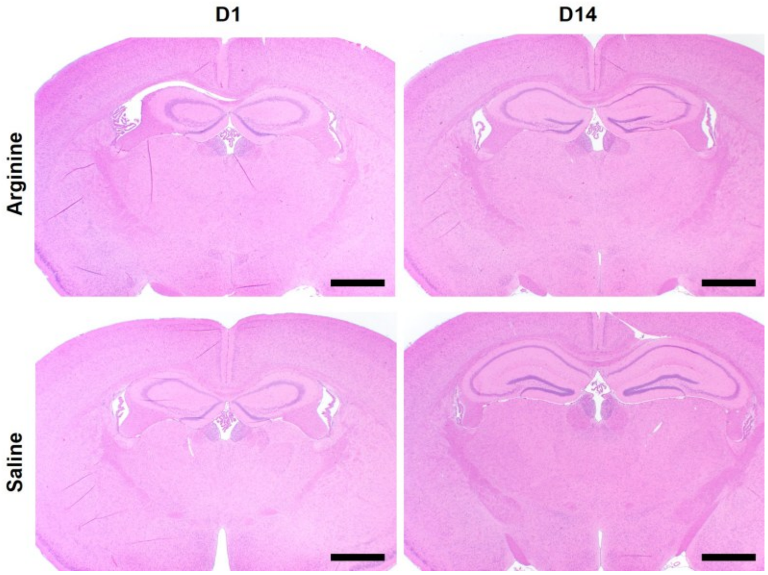


**Fig. S9.** Histology images of H&E-stained brain tissue (cerebrum) sections from experimental mouse groups at 24 h (D1) and 2 weeks (D14) following topical application of the 2 M Arg·HCl solution to the abdominal area (n = 4 per time point). Mice treated with saline alone were used as the control group (n = 4 per time point). Scale bars are 1 mm.

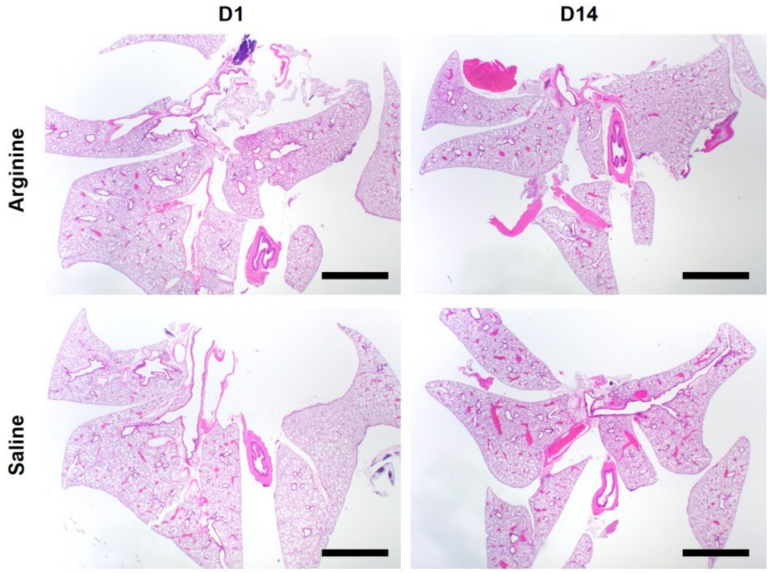


**Fig. S10.** Histology images of H&E-stained lung tissue sections from experimental mouse groups at 24 h (D1) and 2 weeks (D14) following topical application of the 2 M Arg·HCl solution to the abdominal area (n = 4 per time point). Mice treated with saline alone were used as the control group (n = 4 per time point). Scale bars are 2 mm.

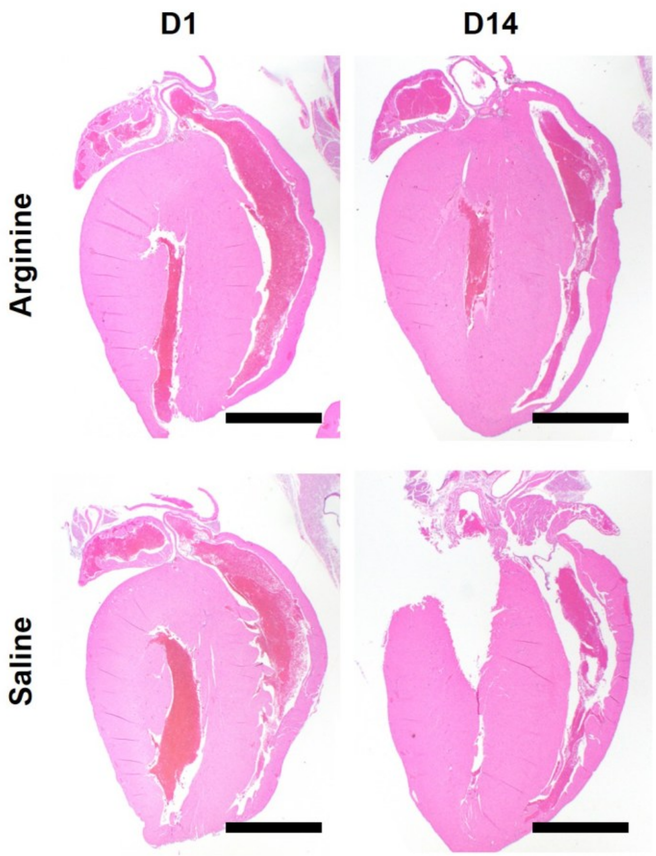


**Fig. S11.** Histology images of H&E-stained heart tissue sections from experimental mouse groups at 24 h (D1) and 2 weeks (D14) following topical application of the 2 M Arg·HCl solution to the abdominal area (n = 4 per time point). Mice treated with saline alone were used as the control group (n = 4 per time point). Scale bars are 2 mm.

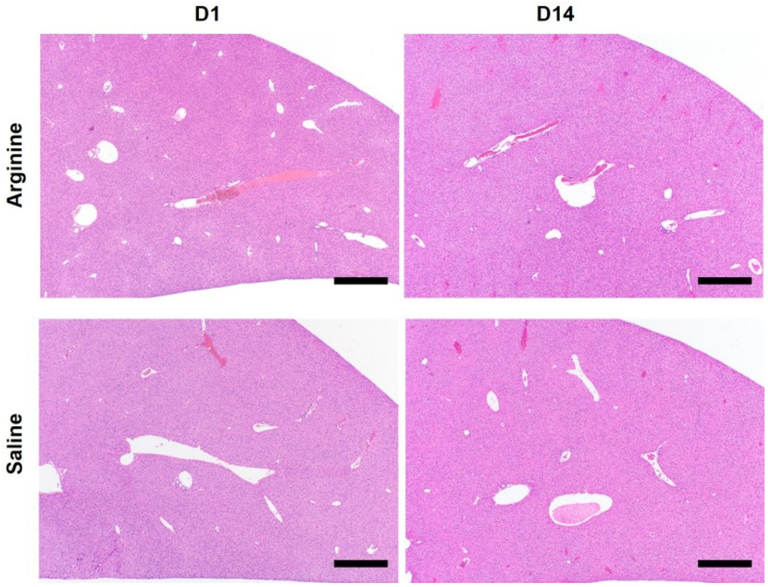


**Fig. S12.** Histology images of H&E-stained liver tissue sections from experimental mouse groups at 24 h (D1) and 2 weeks (D14) following topical application of the 2 M Arg·HCl solution to the abdominal area (n = 4 per time point). Mice treated with saline alone were used as the control group (n = 4 per time point). Scale bars are 500 μm.

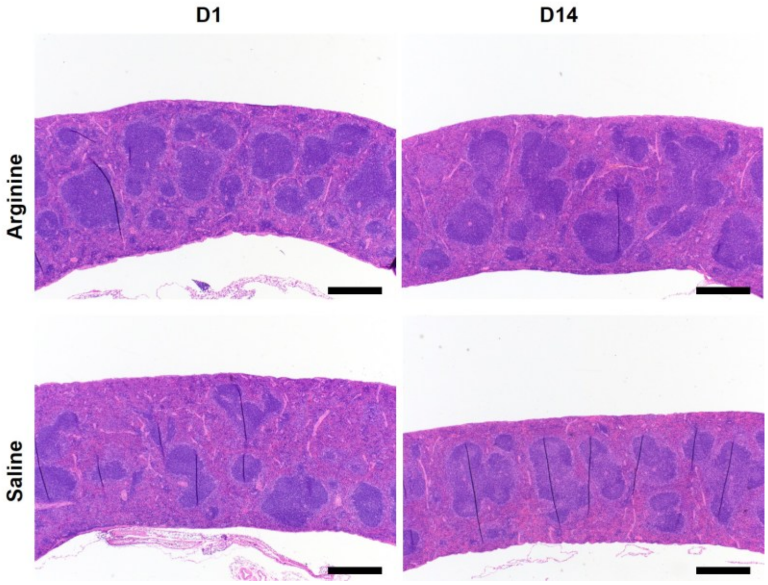


**Fig. S13.** Histology images of H&E-stained spleen tissue sections from experimental mouse groups at 24 h (D1) and 2 weeks (D14) following topical application of the 2 M Arg·HCl solution to the abdominal area (n = 4 per time point). Mice treated with saline alone were used as the control group (n = 4 per time point). Scale bars are 500 μm.

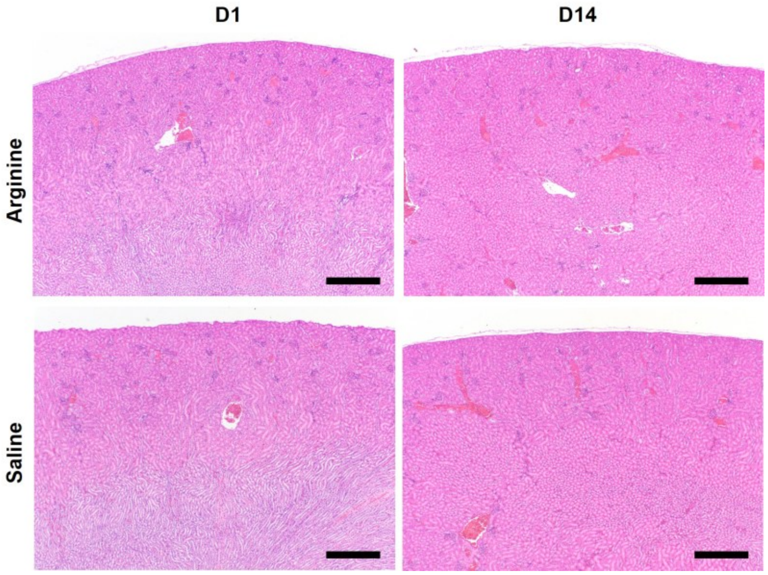


**Fig. S14.** Histology images of H&E-stained kidney tissue sections from experimental mouse groups at 24 h (D1) and 2 weeks (D14) following topical application of the 2 M Arg·HCl solution to the abdominal area (n = 4 per time point). Mice treated with saline alone were used as the control group (n = 4 per time point). Scale bars are 500 μm.

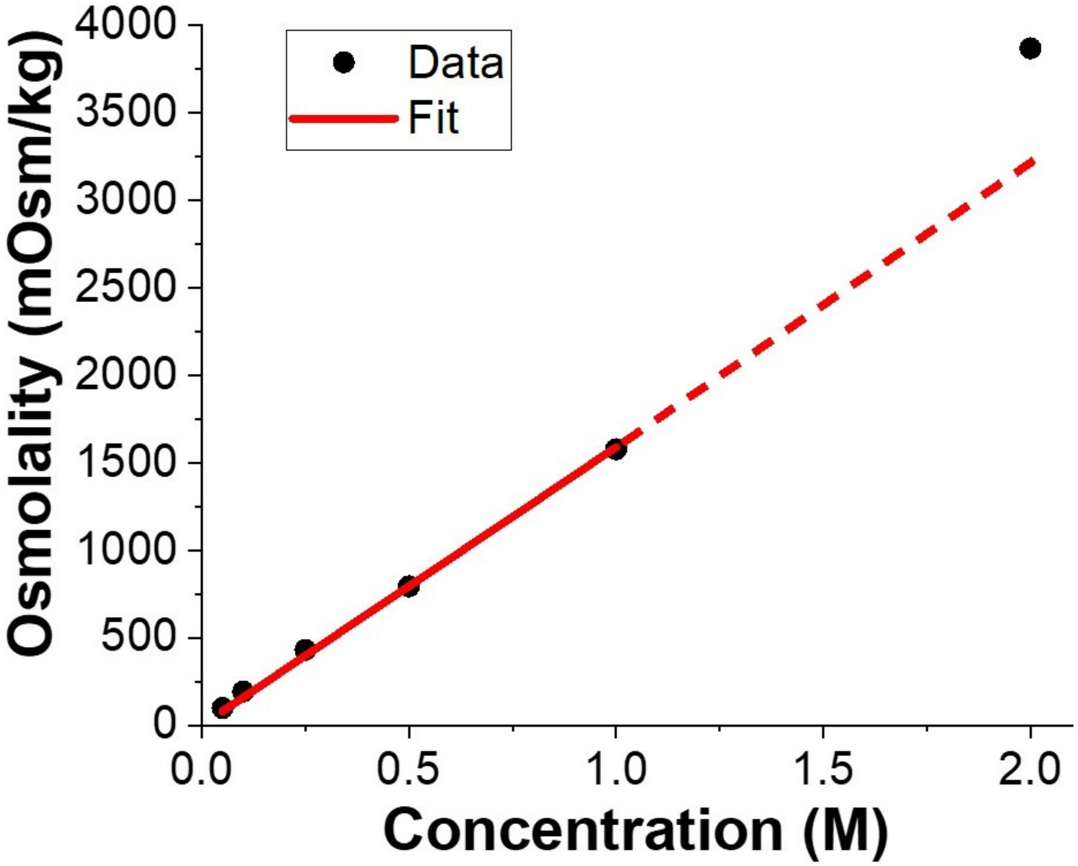


**Fig. S15.** Osmolality plotted as a function of the concentration of Arg·HCl in an aqueous solution. Linear regression of measurements up to 1.0 M reveals a slope of $1.59\times10^3$ mOsm $kg^{-1}$ $M^{-1}$. In contrast, the measured osmolality of 2.0 M Arg·HCl exhibited a supralinear deviation from ideal linearity, likely due to the strong propensity of arginine molecules to form hydrogen bonds with water. This interaction has been reported to reduce the thermodynamic activity of water, resulting in an elevated apparent osmolality.[1]

**Table S1.** Skin histology scores for individual parameters. For scoring of epidermis/dermis changes,1 denotes ≤ 25% of evaluated section, 2 denotes 25-49% of evaluated section, 3 denotes 50-74% of evaluated section, and 4 denotes ≥ 75% of evaluated section. For scoring of panniculitis and myositis, 1 = yes and 0 = no. Lower scores indicate better skin recovery. The similar scoring in the arginine-treated and saline-treated groups suggests that these skin changes result from mechanical rubbing rather than from the clearing agent, arginine hydrochloride, itself.

| Days Post-Treatment | Topically Application | Epidermis/ Dermis | Panniculitis? Y/N | Myositis? Y /N | Final score |
|---|---|---|---|---|---|
| D1 | 2 M Arg·HCl | 1 | 1 | 0 | 2 |
| | | 2 | 1 | 0 | 3 |
| | | 2 | 1 | 0 | 3 |
| | | 3 | 1 | 0 | 4 |
| | 1x PBS | 1 | 1 | 0 | 2 |
| | | 2 | 1 | 0 | 3 |
| | | 3 | 1 | 0 | 4 |
| | | 3 | 1 | 0 | 4 |
| D14 | 2 M Arg·HCl | 1 | 1 | 0 | 2 |
| | | 2 | 1 | 0 | 3 |
| | | 1 | 1 | 1 | 3 |
| | | 1 | 1 | 0 | 2 |
| | 1x PBS | 2 | 0 | 0 | 2 |
| | | 1 | 1 | 0 | 2 |
| | | 1 | 0 | 0 | 1 |

| | | | | | |
|---|---|---|---|---|---|
| | | 1 | 0 | 0 | 1 |

### Supplementary References

(1) Arbelaez-Camargo, D.; Roig-Carreras, M.; García-Montoya, E.; Pérez-Lozano, P.; Miñarro-Carmona, M.; Ticó-Grau, J. R.; Suñé-Negre, J. M. Osmolality Predictive Models of Different Polymers as Tools in Parenteral and Ophthalmic Formulation Development. *Int. J. Pharm.* **2018**, *543* (1-2), 190–200.
